# Meningeal lymphatic drainage promotes T cell responses against *Toxoplasma gondii* but is dispensable for parasite control in the brain

**DOI:** 10.1101/2022.06.02.494581

**Authors:** Michael A. Kovacs, Maureen N. Cowan, Isaac W. Babcock, Lydia A. Sibley, Katherine Still, Samantha J. Batista, Sydney A. Labuzan, Ish Sethi, Tajie H. Harris

**Author notes:** Corresponding Author and Lead Contact: Tajie H. Harris, 409 Lane Road, MR-4 Room 6148, Charlottesville, VA 22908. This work was funded by National Institute of Health grants R01NS112516, R01NS091067, and R56NS106028 to T.H.H.; F30AI154740, 5T32AI007496 and 5T32GM007267 to M.A.K.; T32AI007496 to M.N.C. and I.W.B.; T32GM008328 to K.S.; and T32AI007046 to S.J.B. This work was also funded by the University of Virginia Pinn Scholars Award.

## Abstract

The discovery of meningeal lymphatic vessels that drain the central nervous system (CNS) has prompted new insights into how neuroinflammation develops. In this study, we examined how T cell responses against CNS-derived antigen develop in the context of infection. We found that meningeal lymphatic drainage promotes CD4^+^ and CD8^+^ T cell responses against the neurotropic parasite *Toxoplasma gondii*, and we discovered changes in the antigen-presenting cell compartment of the dural meninges that potentially support this process. Indeed, compared to uninfected controls, mice chronically infected with *T. gondii* displayed a ten-fold increase in the total number of dendritic cells in the dural meninges. These cells upregulated MHC class II, CD80, and CD86 expression, sampled cerebrospinal fluid-derived protein, and were detected within meningeal lymphatic vessels in greater numbers during infection. Disrupting meningeal lymphatic drainage via ligation surgery resulted in reduced dendritic cell number and maturation in the deep cervical lymph nodes and impaired CD4^+^ and CD8^+^ T cell activation, proliferation, and IFN-γ production at this site. Surprisingly, parasite-specific T cell responses in the brain remained intact following ligation, which may be due to activation of T cells at alternative sites during chronic infection, including lymph nodes that drain non-CNS tissue. Collectively, our work reveals that CNS lymphatic drainage supports the development of peripheral T cell responses against *T. gondii* but is nonetheless dispensable for host protection of the brain.

## INTRODUCTION

Although the presence of T cells in the central nervous system (CNS) can be pathological, T cells play an essential role in orchestrating host-protective immune responses in the CNS during infection [1, 2]. At present, the biological mechanisms that drive the formation of T cell responses in the CNS remain poorly defined. One well-studied brain-tropic pathogen is *Toxoplasma gondii*, an intracellular protozoan parasite that causes chronic, lifelong infection in a wide range of mammalian hosts, including humans [3, 4]. Although the course of infection is typically asymptomatic in humans, severe neurologic disease can manifest in immunocompromised individuals, including HIV/AIDS patients [5]. Consistent with this susceptibility, experimental studies have demonstrated that mice deficient in T cell responses are unable to protect the host from fatal disease [6]. For this reason, murine infection with *T. gondii* is a useful model for investigating how T cell responses develop in the CNS.

At homeostasis, the immunologically quiescent brain harbors very few T cells, but in response to *T. gondii* infection, parasite-specific T cells are actively and continuously recruited to the brain [7]. The process is tightly regulated, as circulating T cells that have been activated in the periphery can only enter and persist within the brain when their cognate antigen is expressed in the brain [8, 9]. Disrupting entry of newly activated T cells into the brain leads to impaired parasite restriction, suggesting that ongoing T cell stimulation in the periphery is essential [7, 9]. However, because the brain parenchyma does not harbor lymphatic vessels, it remains poorly understood how peripheral T cells sense microbial antigen expressed in the brain.

In other organs, lymphatic vessels serve as conduits for the transport of tissue-derived antigen and dendritic cells to lymph nodes, where naïve and memory T cells are optimally positioned for detection of their cognate antigen [10, 11]. The recent discovery of functional lymphatic vessels in the dura mater layer of meninges has prompted a significant reconsideration of how the CNS engages the peripheral immune system [12, 13]. Meningeal lymphatic vessels are observed in rodents, primates, and humans [14, 15], and in experimental models of brain cancer and autoimmunity these vessels have been shown to play an integral role in regulating T cell responses in the CNS [16, 17].

Mouse studies have demonstrated that meningeal lymphatic vessels convey macromolecules and immune cells from the meninges and cerebrospinal fluid (CSF) to the deep cervical lymph nodes [17]. The drainage of CSF components by the meningeal lymphatic system is significant because CSF freely exchanges with brain interstitial fluid (ISF) and acts as a sink for the clearance of macromolecules from the brain parenchyma [18, 19]. When model antigens like ovalbumin (OVA) are injected into the brain, these molecules are directed from the ISF into the CSF by glymphatic flow [20], and when injected into the brain or expressed ectopically by brain-resident cells, these proteins have the potential to be presented to T cells in the deep cervical lymph nodes [21, 22].

In a murine model of acute brain infection, it was shown that meningeal lymphatic drainage contributes to the clearance of Japanese encephalitis virus from the brain and promotes host survival [23]. However, questions remain regarding how the meningeal lymphatic system affects T cell responses against brain-derived microbial antigen and the specific role that meningeal lymphatic drainage plays during chronic brain infection. In this study, we report an expansion of the antigen-presenting cell compartment in the dural meninges following infection with *T. gondii*, and we find that these cells sample CSF-derived protein and access meningeal lymphatic vessels in greater numbers during infection. We show that meningeal lymphatic drainage is required for dendritic cell responses in the deep cervical lymph nodes and promotes robust parasite-specific T cell responses at this site. However, in contrast to the finding that meningeal lymphatic drainage is host-protective during acute viral infection of the brain [23], we observed that meningeal lymphatic drainage is dispensable for controlling chronic *T. gondii* infection of the brain, with the T cell response in the brain remaining intact following surgical disruption of meningeal lymphatic outflow. Concurrent activation of T cells observed at alternative sites, including lymph nodes that drain non-CNS tissue, potentially explains the durability of the T cell response in the brain. Overall, our findings highlight how the dural meninges mediates cross-talk between the CNS and peripheral immune system during chronic brain infection, while providing further evidence for the role of meningeal lymphatic drainage in supporting T cell responses against brain-derived antigen.

## RESULTS

### Expansion of the dendritic cell population in the dural meninges during chronic brain infection

The dura mater layer of meninges has emerged as an important anatomic site for immune surveillance of the CNS [24, 25]. Soluble brain- and CSF-derived molecules accumulate within the meninges around the dural sinuses and can be captured by local antigen-presenting cells (APCs) [17, 26]. Moreover, in contrast to the brain, which is devoid of lymphatic vessels, the dural meninges develop an extensive network of lymphatic vessels which directly transport soluble brain- and CSF-derived molecules to peripheral lymph nodes [13, 17]. At homeostasis, a small number of APCs in the dural meninges surround these lymphatic vessels and upon stimulation can traffic to the deep cervical lymph nodes [17].

Although dendritic cells accumulate in the brain during *T. gondii* infection [27, 28], there is only limited evidence to suggest that, in the absence of local lymphatic vasculature, these cells are able to migrate out of the brain to transport antigen directly to peripheral lymph nodes [29, 30]. By contrast, professional APCs in the dural meninges are uniquely positioned to sample CNS material and traffic through lymphatic vessels to lymph nodes for T cell activation. Supporting a role for these cells in the immune response to brain infection, we report a striking accumulation of CD11c^+^ cells in the dural meninges of CD11c^YFP^ reporter mice chronically infected with the ME49 strain of *T. gondii* (**Figure 1a-b**). Quantification by flow cytometry revealed a greater than ten-fold increase in the number of dural dendritic cells (CD45^+^TCRβ^-^NK1.1^-^CD19^-^CD11c^hi^MHC II^hi^) in infected compared to naïve C57BL/6 mice (**Figure 1c, Figure 1 Supp 1**). Intriguingly, the expansion of the dendritic cell compartment in the dural meninges occurred independently of direct infection of the dural meninges. Using real-time PCR to quantify parasite burden by genomic DNA copy number, *T. gondii* was detected in the brain but not in the dural meninges at 6 weeks post-infection (wpi) (**Figure 1d**). These data reinforce an emerging view that immune cells localized to border tissues of the CNS, including the dural meninges, are poised to respond to pathologic disturbances in the CNS [24-26, 31, 32].

**Figure 1.**
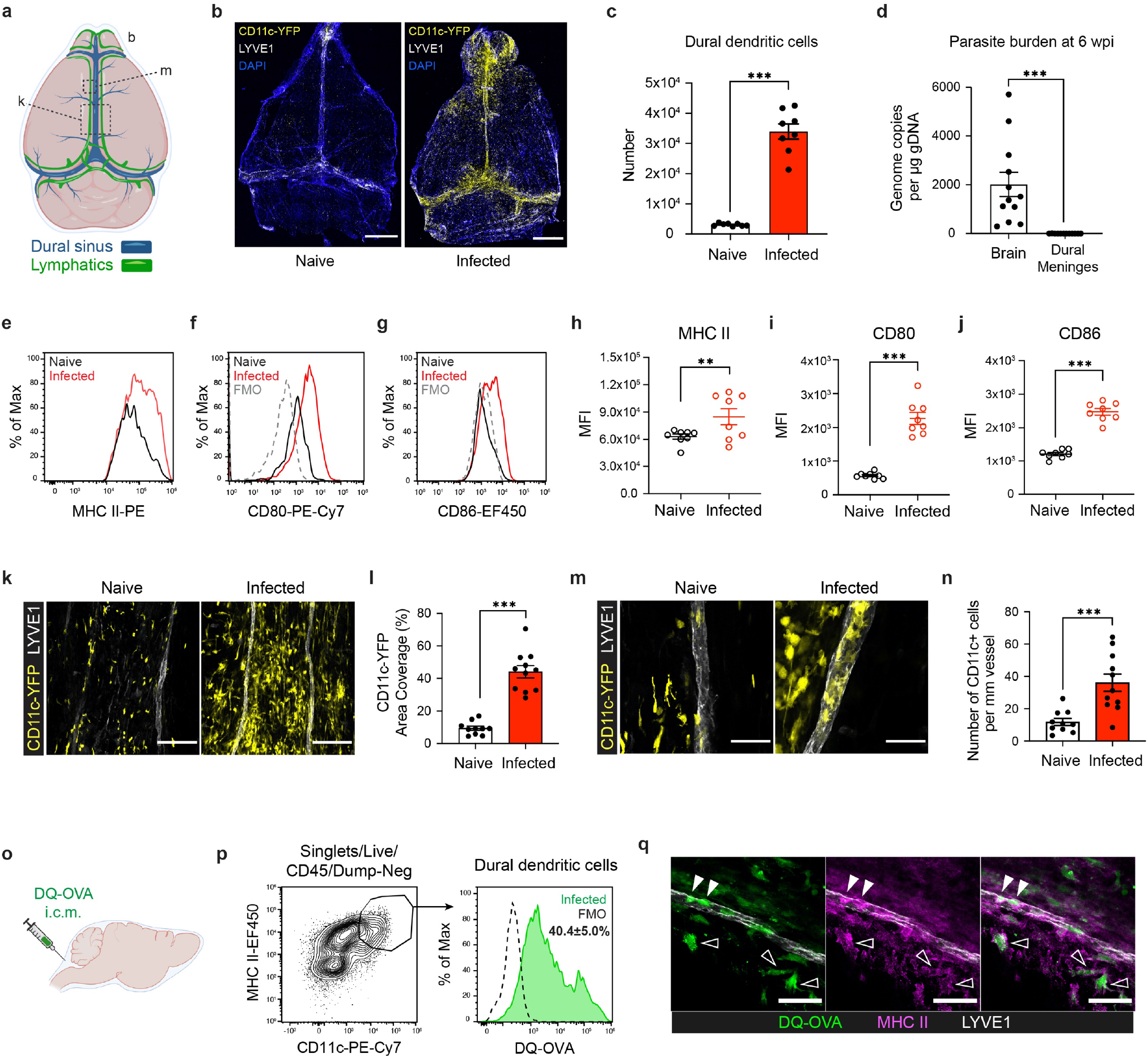
Activated dendritic cells accumulate in the dural meninges and sample CSF-derived protein during chronic brain infection. Mice were infected with 10 cysts of the ME49 strain of *T. gondii* intraperitoneally (i.p.) and analyzed at 6 weeks post-infection. **a**, Schematic diagram of the meningeal lymphatic system underlying the dorsal aspect of the skull. **b**, Confocal microscopy was used to image whole-mount dural meninges dissected from the skullcaps of naïve or chronically infected CD11c^YFP^ reporter mice. Images are representative of three independent experiments. Scale bar, 2000 μm. **c**, Quantification by flow cytometry of total dendritic cell number (CD45^+^TCRβ^-^NK1.1^-^CD19^-^CD11c^hi^MHC II^hi^) in the dural meninges of naïve or chronically infected C57BL/6 mice. Data compiled from two experiments. **d**, Quantification of *T. gondii* gDNA in brain and dural meninges by real-time PCR. Data compiled from three experiments. **e-g**, Representative flow histograms showing MHC class II **(e)**, CD80 **(f)**, and CD86 **(g)** expression by dendritic cells in the dural meninges of naïve or chronically infected C57BL/6 mice. **h-j**, Quantification of the geometric mean fluorescence intensity (MFI) of MHC class II **(h)**, CD80 **(i)**, and CD86 **(j)** expressed by dendritic cells in the dural meninges. Data compiled from two experiments. **k**, Representative images of CD11c-YFP^+^ cells (yellow) in the region surrounding the dural sinuses. Scale bar, 150 μm. **l**, Quantification of area coverage by CD11c- YFP^+^ cells in the region surrounding the dural sinuses. Data compiled from three experiments. **m- n**, Representative images **(m)** and quantification **(n)** of CD11c-YFP^+^ cells (yellow) present within LYVE1^+^ meningeal lymphatic vessels (white). Data compiled from three experiments. Scale bar, 50 μm. **o**, Schematic diagram illustrating intra-cisterna magna (i.c.m.) injection of DQ-OVA into the CSF. **p**, Representative flow histogram for fluorescent emission of proteolytically cleaved DQ- OVA in dural dendritic cells 5 h after i.c.m. injection into chronically infected mice. The average frequency of DQ-OVA^+^ dendritic cells (mean value ± s.e.m.) was calculated from three pooled experiments. **q**, Representative images of MHC class II-expressing antigen-presenting cells (magenta) co-labeling with fluorescent DQ-OVA cleavage products (green) in the vicinity of (open arrows) or present within (closed arrows) meningeal lymphatic vessels (LYVE1^+^, white). Three independent experiments were performed. Scale bar, 50 μm. For pooled experiments (**c-d**, **h-j**, **l**, **n**), data are represented as mean values ± s.e.m. and statistical significance was measured using randomized block ANOVA (two-way), with *p* < 0.01 (**) and *p* < 0.001 (***).

Dendritic cells can undergo maturation in response to variety of inflammatory stimuli to become more efficient APCs [33]. In response to chronic brain infection, dendritic cells in the dural meninges displayed an increase in surface expression of MHC class II and the co-stimulatory molecules CD80 and CD86, suggesting that chronic brain infection promotes maturation of APCs in the dural meninges (**Figure 1e-j**). CD11c^+^ cells localized preferentially around the dural sinuses near meningeal lymphatic vessels (**Figure 1b, 1k**), increasing local area coverage from 9% in naïve mice to 44% in chronically infected mice (**Figure 1l**). This pattern of distribution is interesting because prior studies have demonstrated that protein tracers injected into the brain or CSF accumulate in this same region around the dural sinuses [17, 26], suggesting that these cells may be poised to sample CNS-derived protein during infection.

Notably, three times the number of CD11c^+^ cells were present in LYVE1^+^ meningeal lymphatic vessels in infected mice (**Figure 1m-n**), suggesting that dural APCs are trafficking in response to brain infection. Because activated dendritic cells injected into the CSF have the capacity to enter meningeal lymphatic vessels and drain to the deep cervical lymph nodes [17], it is possible that some of these dendritic cells originated in the CSF. Therefore, we performed flow cytometry on CSF pooled from 4-5 naïve or chronically infected mice and determined that, in fact, a population of dendritic cells not present in naïve mice emerged in the CSF of brain-infected mice (**Figure 1 Supp 2a-b**), expressing high levels of CD80 and CD86 (**Figure 1 Supp 2c-d**). Interestingly, the concentration of CSF-borne dendritic cells was consistent with that reported by a study of human CSF samples isolated from patients with Lyme neuroborreliosis [34].

We next sought to determine whether APCs trafficking in the meningeal lymphatic vessels during *T. gondii* brain infection are able to transport CSF-derived protein. CSF has been shown to function as a sink for brain-derived macromolecules [18, 19] and would therefore represent a potential mechanism by which APCs in the dural meninges could sample brain-derived antigen during infection. To address this question, soluble DQ-OVA was injected into the CSF of chronically infected mice via intra-cisterna magna (i.c.m.) injection (**Figure 1o**). DQ-OVA is a self-quenched conjugate of ovalbumin that only emits fluorescence after proteolytic cleavage and has been used to examine uptake and processing of soluble antigen by dendritic cells [21, 35, 36]. Interestingly, by flow cytometry it was observed that 40% of dendritic cells in the dural meninges harbored DQ-OVA cleavage products five hours after i.c.m. injection (**Figure 1p**). Moreover, immunostaining of whole-mount meninges revealed that DQ-OVA^+^ cells were present within meningeal lymphatic vessels, implying that dural APCs can capture and transport CSF-borne antigen during chronic brain infection (**Figure 1q**).

All together, these findings highlight significant changes in the APC compartment of the dural meninges in response to *T. gondii* brain infection. Dendritic cells at this site increased in number, displayed a more activated phenotype, were able to sample CSF-borne protein, and accumulated within meningeal lymphatic vessels.

### T cell responses in the deep cervical lymph nodes peak following progression to chronic brain infection

Initial studies in mice, and later in humans, have demonstrated that meningeal lymphatic vessels drain directly to the deep cervical lymph nodes (DCLN) [15, 17]. Even before the immunologic function of the meningeal lymphatic system started coming into focus, these lymph nodes were strongly implicated in regulating CNS immunity. For example, injection of antigen into rodent brains elicited strong humoral responses in the DCLNs [37], and surgical excision of the DCLNs contributed to a reduction in severity of experimental autoimmune encephalomyelitis [38, 39]. To date, few studies have examined the contribution of the deep cervical lymph nodes to immunity against CNS pathogens. To address the role of the DCLNs during *T. gondii* infection, we first confirmed that during chronic infection CSF-derived protein could be sampled by dendritic cells present within these lymph nodes. Upon i.c.m. injection of DQ-OVA into chronically infected mice, 3.3% of dendritic cells in the DCLNs were found to harbor digested protein products after 5 hours (**Figure 2a-b**). By contrast, processed DQ-OVA was not detected in the inguinal lymph nodes (ILN), which drain the hindlimb (**Figure 2a-b**). Because soluble protein can drain directly to lymph nodes or can be transported by migratory dendritic cells, we were unable to distinguish what fraction of DQ-OVA^+^ dendritic cells in the DCLNs had endocytosed protein locally and what fraction had engulfed protein in the dural meninges before trafficking to the DCLNs (**Figure 1q**). Whatever their origin, dendritic cells present in the deep cervical lymph nodes efficiently captured CSF-derived protein during *T. gondii* infection.

**Figure 2.**
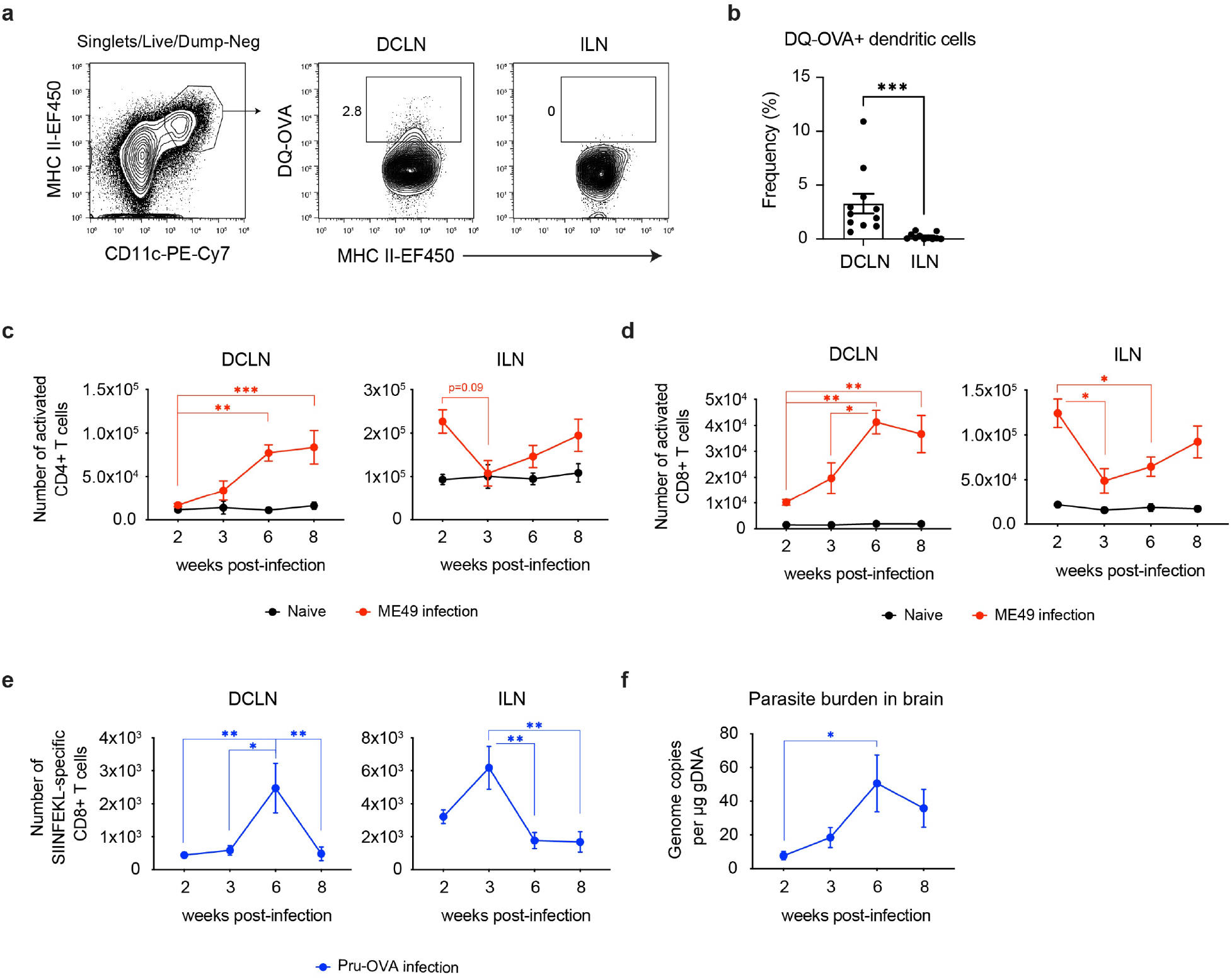
Expansion of T cells in the deep cervical lymph nodes occurs primarily during the chronic stage of infection and tracks with parasite burden in the brain. **a-b**, DQ-OVA was injected into the CSF of chronically infected mice by i.c.m. injection and fluorescent emission of proteolytically cleaved DQ-OVA was measured in CD11c^hi^MHC II^hi^ dendritic cells of the deep cervical lymph nodes (DCLNs) and inguinal lymph nodes (ILNs) 5 h later by flow cytometry. **(a)** Representative contour plots of DQ-OVA^+^ dendritic cells in the DCLNs or ILNs at 6 wpi. CD11c^hi^MHC II^hi^ cells were pre-gated on singlets/live/TCRβ^-^/NK1.1^-^/CD19^-^. **(b)** Quantification of frequency of DQ-OVA^+^ dendritic cells in the DCLNs or ILNs at 6 wpi. Data are compiled from three experiments and are represented as mean values ± s.e.m. Statistical significance was measured using randomized block ANOVA (two-way), with *p* < 0.001 (***). **c-d**, C57BL/6 mice were infected i.p. with 10 cysts of the ME49 strain of *T. gondii* and total number of activated CD4^+^ T cells **(c)** or activated CD8^+^ T cells **(d)** in the DCLNs or ILNs was quantified at multiple time points over the course of acute and chronic infection (red dots). The steady-state number of activated CD4^+^ and CD8^+^ T cells in the different lymph node compartments was measured in naïve mice at corresponding time points (black dots). Activated T cells displayed a CD44^hi^CD62L^lo^ phenotype. Data are compiled from three experiments. **e-f**, C57BL/6 mice were infected i.p. with 1,000 tachyzoites of the Pru-OVA strain of *T. gondii*. **(e)** Total number of SIINFEKL-specific CD8^+^ T cells in the DCLNs or ILNs was quantified at multiple time points over the course of acute and chronic infection using tetramer reagent. **(f)** Quantification of *T. gondii* gDNA in brain at corresponding time points by real-time PCR. Data are compiled from two experiments. For time course experiments (**c-f**), data are represented as mean values ± s.e.m. and statistical significance of differences across time points in infected mice was measured using one-way ANOVA with post-hoc Tukey multiple comparison testing. *p* < 0.05 (*), *p* < 0.01 (**), and *p* < 0.001 (***).

Based on these results, we performed a time course experiment to examine the kinetics of T cell responses in the deep cervical lymph nodes. Importantly, infection with type II strains of the parasite, including ME49, involves two distinct stages [40]. During acute infection, which lasts two to three weeks, *T. gondii* rapidly disseminates throughout the host’s peripheral tissues as fast-replicating tachyzoites, before being cleared by a robust IFN-γ response. During chronic infection, the parasite converts to slow-replicating bradyzoites, which persist in the brain and, to a limited extent, skeletal muscle as semi-dormant cysts. When C57BL/6 mice were infected with the ME49 strain of *T. gondii*, activated (CD44^hi^CD62L^lo^) CD4^+^ and CD8^+^ T cell number in the hindlimb-draining ILNs was greatest at 2 wpi during acute infection (**Figure 2c-d, Figure 2 Supp 1a**), consistent with the broad distribution of parasite in peripheral tissues at this time point. Conversely, only a small number of activated CD4^+^ and CD8^+^ T cells was detected in the DCLNs at 2 wpi (**Figure 2c-d**). Expansion of CD4^+^ and CD8^+^ T cells in the DCLNs became more pronounced after progression to the chronic stage of infection, with the peak in number of activated CD4^+^ and CD8^+^ T cells occurring at 6-8 wpi (**Figure 2c-d**). These results indicate that the deep cervical lymph nodes play a distinct role from other lymph nodes in supporting T cell responses against *T. gondii* specifically during chronic infection.

To better understand the stage-dependent activation of T cells in the DCLNs, we used MHC class I tetramer to track a parasite epitope (SIINFEKL)-specific population of CD8^+^ T cells generated in response to infection with Pru-OVA, a recombinant type II strain of *T. gondii* engineered to express a secreted, truncated form of the model antigen ovalbumin [41] (**Figure 2 Supp 1b**). Consistent with the polyclonal CD8^+^ T cell response observed in response to ME49 infection (**Figure 2d**), the peak SIINFKEKL-specific CD8^+^ T cell response in the hindlimb-draining ILNs occurred during acute infection (2-3 wpi), when very few parasite-specific T cells were detectable in the DCLNs (**Figure 2e**). By contrast, expansion of parasite-specific CD8^+^ T cells in the DCLNs was greatest during chronic infection, increasing significantly between 3 wpi and 6 wpi and tracking closely with parasite burden in the brain, which similarly peaked at 6 wpi (**Figure 2e-f**).

These results indicate that the magnitude of T cell responses in different lymph node compartments change as the tissue distribution of parasite evolves over the course of infection, with T cell responses in the deep cervical lymph nodes corresponding closely with infection of the brain. These data suggest that the deep cervical lymph nodes play a specific role in mediating T cell responses against brain-derived antigen during *T. gondii* infection.

### Meningeal lymphatic drainage promotes peripheral T cell responses against *T. gondii*

To directly test the function of meningeal lymphatic drainage during *T. gondii* brain infection, we surgically ligated the collecting vessels afferent to the deep cervical lymph nodes. This approach has been used to test the function of meningeal lymphatic drainage in experimental models of multiple sclerosis, glioblastoma, and acute viral infection of the brain [16, 17, 23]. We performed ligation at 3 wpi, the onset of chronic infection, and analyzed mice 3 weeks later (**Figure 3a**). Unless otherwise stated, experiments were performed using the ME49 strain of *T. gondii*. Tracer studies using Evans blue or Alexa Fluor 594-conjugated ovalbumin (OVA-AF594) confirmed the efficacy of the procedure and demonstrated a greater than 95% reduction in meningeal lymphatic outflow of CSF-derived components in ligated animals (**Figure 3 Supp 1a-b**).

**Figure 3.**
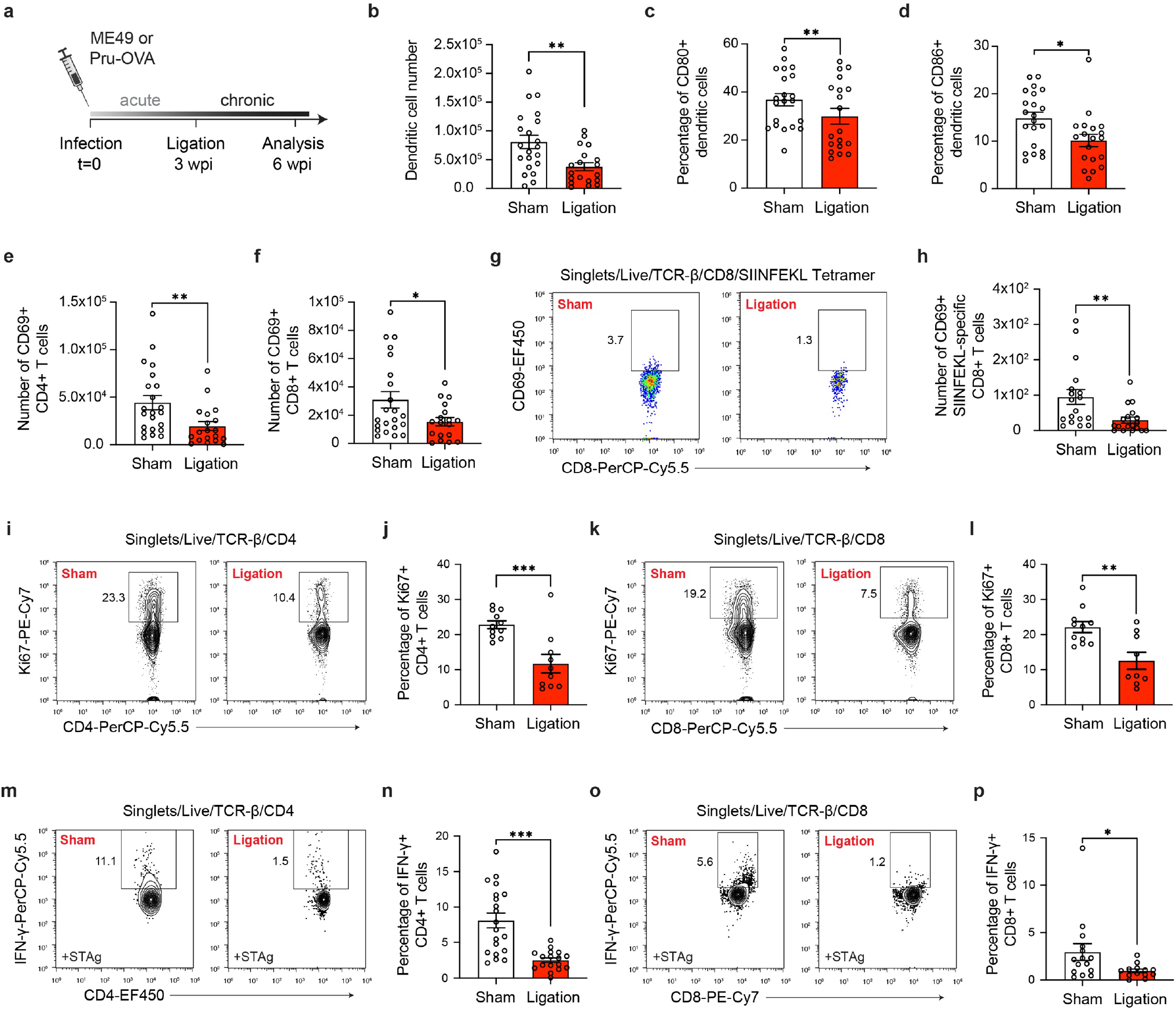
Restricting meningeal lymphatic drainage disrupts T cell activation, proliferation, and cytokine production in the deep cervical lymph nodes. **a**, Experimental design for ligation studies in C57BL/6 mice. **b-d**, Quantification by flow cytometry of the number of dendritic cells observed in the DCLNs three weeks after ligation or sham surgery **(b)** and the frequency of these cells expressing CD80 **(c)** or CD86 **(d)**. Data are compiled from four experiments. **e-f**, Quantification by flow cytometry of the number of CD69-expressing CD4^+^ T cells **(e)** or CD8^+^ T cells **(f)** in the DCLNs three weeks after ligation or sham surgery. Data are compiled from four experiments. **g-h**, Representative dot plots **(g)** and total number **(h)** of CD69-expressing SIINFEKL-specific CD8^+^ T cells in the DCLNs three weeks after ligation or sham surgery of mice infected with Pru-OVA. Data are compiled from four experiments. **i-j**, Representative contour plots **(i)** and average frequency **(j)** of Ki67-expressing CD4^+^ T cells in the DCLNs three weeks after ligation or sham surgery. Data are compiled from two experiments. **k-l**, Representative contour plots **(k)** and average frequency **(l)** of Ki67-expressing CD8^+^ T cells in the DCLNs three weeks after ligation or sham surgery. Data are compiled from two experiments. **m-p**, Intracellular staining of IFN-γ was performed on cells isolated from the DCLNs of ligated or sham-operated mice 24 h after *ex vivo* restimulation with soluble tachyzoite antigen (STAg). Representative contour plots **(m, o)** and average frequency **(n, p)** of CD4^+^ or CD8^+^ T cells that expressed IFN-γ after STAg restimulation are shown. Mice used throughout these experiments were infected with the ME49 strain of *T. gondii*, with the exception of **g** and **h** (as indicated). Data are represented as mean values ± s.e.m. and statistical significance was measured using randomized block ANOVA (two-way), with *p* < 0.05 (*), *p* < 0.01 (**), and *p* < 0.001 (***).

We first examined the effect of meningeal lymphatic drainage on dendritic cell responses in the deep cervical lymph nodes by flow cytometry. Consistent with our imaging data showing enhanced dendritic cell trafficking within meningeal lymphatic vessels during brain infection (**Figure 1m- n**), restricting drainage by ligation reduced dendritic cell number in the DCLNs by nearly 50% (**Figure 3b**). These data suggest that dendritic cells originating in the dural meninges or CSF contribute significantly to the pool of APCs in the DCLNs. Furthermore, a decreased percentage of dendritic cells present in the DCLNs after ligation expressed CD80 and CD86 on their surface, suggesting that in response to brain infection meningeal lymphatic outflow helps maintain a population of mature dendritic cells at this site (**Figure 3c-d**).

We next hypothesized that peripheral T cell responses would be significantly impaired by ligation, particularly after observing significant defects in the dendritic cell population. Indeed, restricting meningeal lymphatic outflow led to a striking decrease in the frequency and number of CD4^+^ and CD8^+^ T cells in the DCLNs expressing the early activation marker CD69 (**Figure 3e-f**, **Figure 3 Supp 2a-d**). Similarly, when Pru-OVA was used to infect mice, ligation caused a decrease in expression of CD69 by SIINFEKL-specific CD8^+^ T cells in the DCLNs (**Figure 3g-h**). To assess whether this decrease in activation was associated with reduced T cell proliferation, we measured Ki67 expression by flow cytometry and observed a concomitant decrease in the frequency of proliferative CD4^+^ and CD8^+^ T cells in ligated mice (**Figure 3i-l**).

During *T. gondii* infection, CD4^+^ and CD8^+^ T cells confer protection to the host by producing large amounts of IFN-γ and TNF-α, cytokines that activate intracellular mechanisms of parasite restriction in infected host cells [42, 43]. To determine whether meningeal lymphatic drainage is required for maintaining high-quality peripheral CD4^+^ and CD8^+^ T cell responses, intracellular cytokine staining was performed on cells that had been isolated from the deep cervical lymph nodes of sham-operated or ligated mice and restimulated *ex vivo* for 24 h with soluble tachyzoite antigen (STAg). Remarkably, IFN-γ production was reduced by 69% among CD4^+^ T cells and 60% among CD8^+^ T cells after ligation surgery (**Figure 3m-p**). TNF-α production was similarly impaired in CD4^+^ T cells isolated from the DCLNs of ligated mice (**Figure 3 Supp 3a-b**). IFN-γ and TNF-α production was not detected in unstimulated CD4^+^ and CD8^+^ T cells, suggesting that T cell responses were parasite-specific (data not shown).

From these experiments, we can conclude that meningeal lymphatic drainage contributes to robust parasite-specific CD4^+^ and CD8^+^ T cell responses in the deep cervical lymph nodes during chronic brain infection. Restriction of meningeal lymphatic outflow caused defects in dendritic cell responses and peripheral CD4^+^ and CD8^+^ T cell activation, proliferation, and cytokine production.

### Meningeal lymphatic drainage is dispensable for host protection of the brain

Several recent studies have demonstrated that meningeal lymphatic drainage plays a key role in supporting T cell responses in the brain against locally injected tumor cells [16, 44]. Moreover, it has been shown that meningeal lymphatic drainage promotes host survival during acute brain infection with Japanese encephalitis virus [23]. Based on these studies, we hypothesized that meningeal lymphatic drainage would be required for host-protective T cell responses in the brain against *T. gondii*. Unexpectedly, restricting meningeal lymphatic drainage had no discernible impact on the magnitude or quality of CD4^+^ and CD8^+^ T cell responses in the brain after ligation, despite impairing T cell responses in the deep cervical lymph nodes (**Figure 4a-e**).

**Figure 4:**
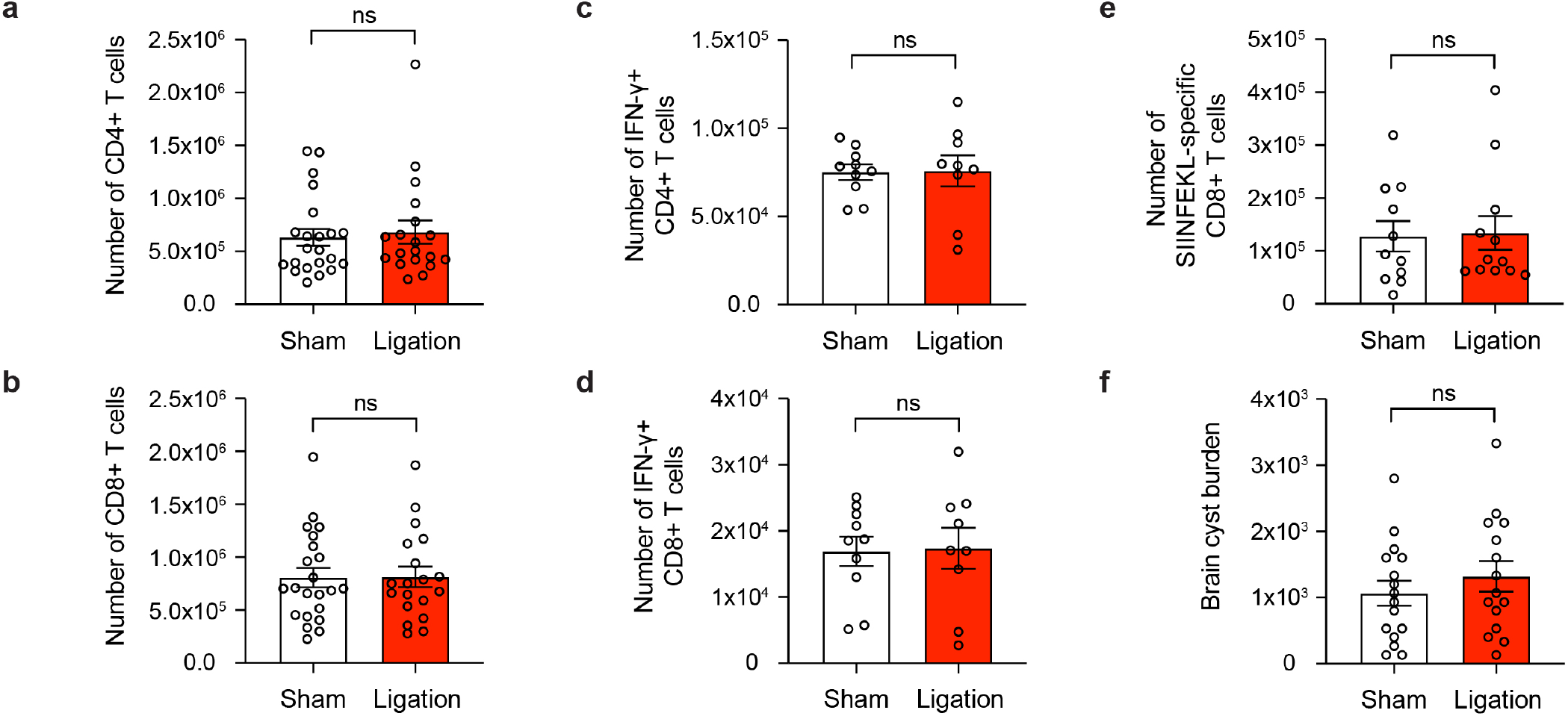
Meningeal lymphatic drainage is dispensable for host-protective T cell responses in the brain. **a-b**, Quantification by flow cytometry of the number of infiltrating CD4^+^ T cells **(a)** and CD8^+^ T cells **(b)** in the brain tissue of ligated or sham-operated mice. Data are compiled from four experiments. **c-d**, Intracellular staining of IFN-γ was performed on mononuclear cells isolated from brains of ligated or sham-operated mice 24 h after *ex vivo* restimulation with soluble tachyzoite antigen (STAg). Total number of CD4^+^ T cells **(c)** and CD8^+^ T cells **(d)** expressing IFN-γ after STAg restimulation are shown. Data are compiled from two experiments. **e**, Quantification by flow cytometry of the number of SIINFEKL-specific CD8^+^ T cells in the brain tissue of ligated or sham-operated mice chronically infected with Pru-OVA. Data are compiled from three experiments. **f**, Quantification of parasite burden by enumeration of tissue cysts in brain homogenate of ligated or sham-operated mice. Data are compiled from three experiments. Data are represented as mean values ± s.e.m. and statistical significance was measured using randomized block ANOVA (two-way), with ns = not significant.

Indeed, the total number of CD4^+^ and CD8^+^ T cells in the brain remained unchanged between sham and ligated mice (**Figure 4a-b**), revealing that sufficient numbers of T cells were recruited to the brain despite impaired parasite-specific T cell responses in the DCLNs. To determine whether there were differences in the quality of T cell responses in the brain, we measured the total number of IFN-γ-producing CD4^+^ and CD8^+^ T cells in the brain following *ex vivo* restimulation with STAg, but again saw no change after ligation (**Figure 4c-d**). Finally, no differences were detected in the total number of SIINFEKL-specific CD8^+^ T cells in the brains of sham-operated or ligated mice chronically infected with Pru-OVA (**Figure 4e**). Because parasite burden was similar between the two groups (**Figure 4f**), it is unlikely that any other mechanism of parasite control became defective in the brain as a consequence of ligation.

These data reveal that meningeal lymphatic drainage is not necessary for maintenance of the host-protective T cell population in the brain during chronic infection with *T. gondii*. These results contrast with recent studies showing that T cell responses against brain tumors and host protection against acute viral infection of the brain are supported by meningeal lymphatic drainage to the deep cervical lymph nodes [16, 23, 44].

### Antigen-dependent stimulation of T cells occurs in CNS- and non-CNS-draining lymph nodes during chronic infection

Because T cell recruitment from the periphery has been demonstrated to be indispensable for T cell responses against *T. gondii* in the brain [7, 9], it is likely that T cell activation during the chronic stage of infection is not limited to the CNS-draining lymph nodes. In order to identify alternative sources of T cells in the periphery, we began by performing a pairwise comparison of T cell proliferation in the deep cervical lymph nodes and inguinal lymph nodes of ME49-infected mice at 6 wpi. A large percentage of CD4^+^ and CD8^+^ T cells expressed Ki67 in the DCLNs at this time point, indicating robust T cell proliferation in CNS-draining lymph nodes during chronic brain infection (**Figure 5a-d**). Interestingly, though, a large population of Ki67-expressing CD4^+^ and CD8^+^ T cells was also detected in the hindlimb-draining ILNs at this time point, albeit at a lower frequency (**Figure 5a-d**). These data suggest that T cell proliferation is enriched in lymph nodes that drain CNS tissue at 6 wpi, when parasite is most abundant in the brain, but also occurs in non-CNS-draining lymph nodes.

**Figure 5:**
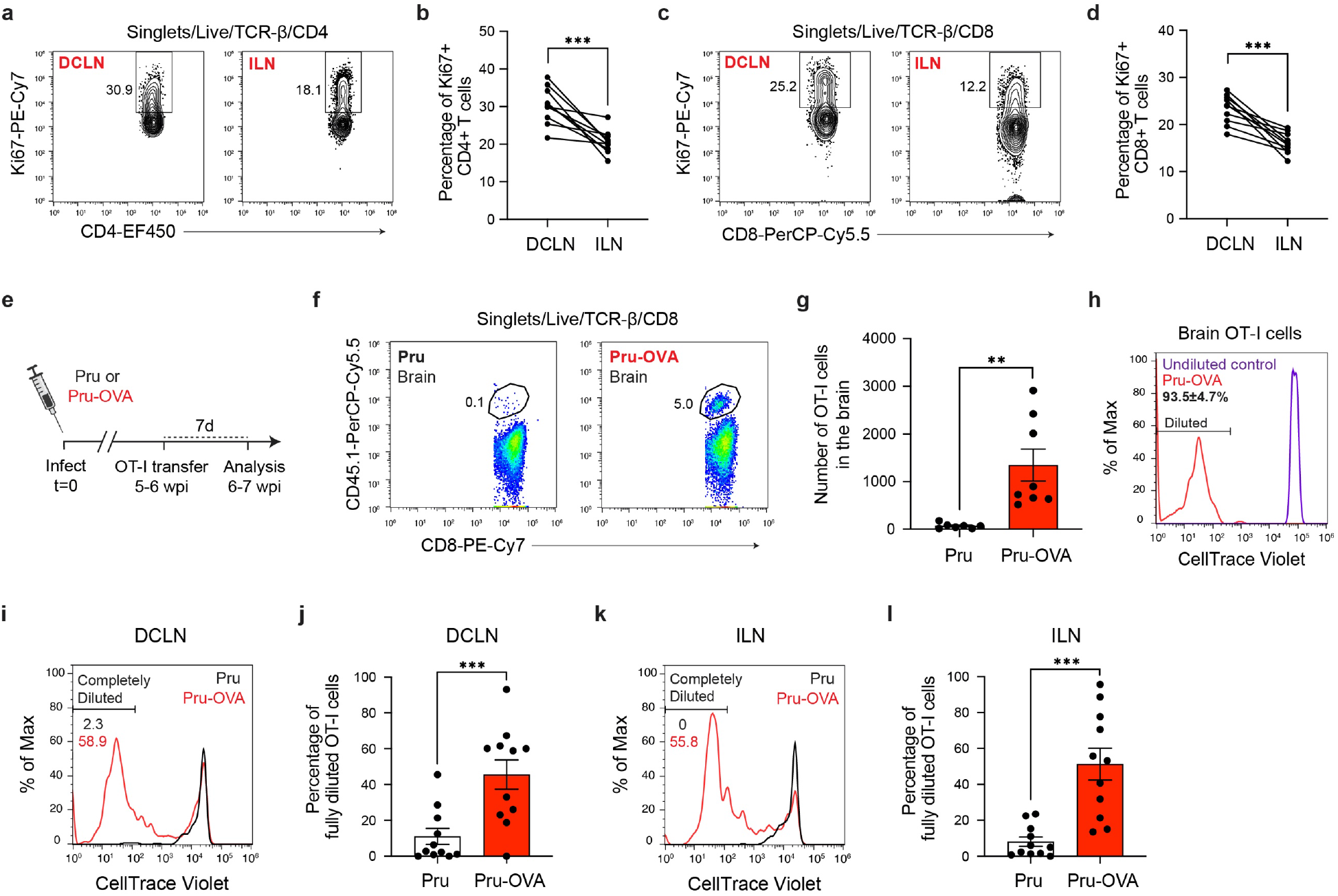
Antigen-dependent proliferation of T cells occurs in CNS- and non-CNS-draining lymph nodes during chronic infection with *T. gondii*. T cell proliferation was measured at 6 wpi in the deep cervical lymph nodes (DCLNs) and a representative group of non-CNS-draining lymph nodes, the inguinal lymph nodes (ILNs). **a-d**, Representative contour plots **(a, c)** and average frequency **(b, d)** of Ki67-expressing CD4^+^ or CD8^+^ T cells in the different lymph node compartments of C57BL/6 mice chronically infected with the ME49 strain of *T. gondii*. Data are compiled from two experiments and statistical significance was measured using a two-tailed paired *t*-test, with *p* < 0.001 (***). **e-l**, C57BL/6 mice were infected with the OVA-secreting Pru strain of *T. gondii* (Pru-OVA) or the control parental Pru (Pru) strain of *T. gondii*. Five to six weeks later, OT-I cells (CD45.1 congenic) labeled with CellTrace Violet were transferred intravenously, and tissues were analyzed seven days post-transfer. **e**, Experimental design for OT-I transfer studies. **f-g**, Representative dot plots **(f)** and total number **(g)** of OT-I cells infiltrating the brain tissue of mice infected with Pru-OVA or parental Pru. Data are compiled from two experiments. **h**, Representative flow histogram showing CellTrace Violet dye dilution among brain-infiltrating OT-I cells of mice chronically infected with Pru-OVA or parental Pru. The average frequency (mean value ± s.e.m.) of fully diluted cells was calculated from two pooled experiments. **i-j**, Representative flow histogram (**i**) and average frequency (**j**) of fully diluted OT-I cells in the DCLNs of mice chronically infected with Pru-OVA or parental Pru. Data are compiled from three experiments. **k-l**, Representative flow histogram (**k**) and average frequency (**l**) of fully diluted OT- I cells in the ILNs of mice chronically infected with Pru-OVA or parental Pru. Data are compiled from three experiments. For all experiments data are represented as mean values ± s.e.m. and for **g, j, and l** statistical significance was measured using randomized block ANOVA (two-way), with *p* < 0.01 (**) and *p* < 0.001 (***).

To more closely examine the locations of antigen stimulation during chronic infection, we adoptively transferred *in vitro*-expanded OVA-specific OT-I cells (CD45.1 congenic) to C57BL/6 mice 5 to 6 weeks after infection with either Pru-OVA or parental strain Pru (**Figure 5e**). We then measured CellTrace Violet dilution in the DCLNs and ILNs one week after transfer. As expected, OT-I cells only accumulated in the brains of mice infected with Pru-OVA (**Figure 5f-g**), and all of the OT-I cells detected in the brains of these mice had fully diluted their CellTrace Violet dye (**Figure 5h**), consistent with the notion that these cells undergo several rounds of proliferation prior to recruitment to the brain.

To understand where peripheral T cells undergo activation and proliferation, we started by measuring CellTrace Violet dilution in the CNS-draining deep cervical lymph nodes. Notably 46% of OT-I cells displayed complete dilution in the DCLNs when mice were infected with Pru-OVA, compared to only 11% of OT-I cells when mice were infected with parental Pru (**Figure 5i-j**). These data indicate that proliferation of SIINFEKL-specific CD8^+^ T cells in CNS-draining lymph nodes is predominantly antigen-dependent, and only a limited degree of proliferation can be attributed to bystander activation. The OT-I cells used for adoptive transfer also expressed GFP under the control of the *Nr4a1* (Nur77) promoter, a reporter system that was originally developed as a tool to track antigen-dependent TCR stimulation [45, 46]. However, we observed that GFP upregulation was equivalent in OT-I cells after infection with either Pru-OVA or parental Pru (**Figure 5 Supp 1a-d**), even though OT-I cells proliferated to a significantly higher degree in the lymph nodes of mice infected with Pru-OVA compared to mice infected with parental Pru (**Figure 5i-l**). To determine whether antigen-dependent T cell stimulation occurred in lymph nodes that drain non-CNS tissue during chronic infection, we measured CellTrace Violet dilution in the inguinal lymph nodes. Intriguingly, the degree of antigen-dependent T cell proliferation at this site was similar to that observed in the deep cervical lymph nodes, as 51% of OT-I cells fully diluted the tracer dye in mice infected with Pru-OVA compared to 8% of OT-I cells in mice infected with parental Pru (**Figure 5k-l**). These data lead us to conclude that antigenic stimulation of T cells occurs concurrently in CNS- and non-CNS-draining lymph nodes during chronic infection. What remains uncertain is the source of antigen eliciting T cell proliferation in the non-CNS-draining lymph nodes during chronic infection, whether residual antigen persisting after acute systemic infection or antigen draining from skeletal muscle, a tissue in which *T. gondii* also encysts.

All together, these experiments reveal that both CNS-draining lymph nodes and non-CNS-draining lymph nodes are sources of newly activated parasite-specific T cells during the chronic stage of *T. gondii* infection. Ongoing proliferation of T cells in non-CNS-draining lymph nodes, such as the inguinal lymph nodes, may explain the durability of T cell responses in the brain when T cell responses in the deep cervical lymph nodes become impaired.

## DISCUSSION

Due to the unique anatomic organization of the CNS [19, 47], notably the absence of lymphatic vessels in the brain parenchyma, it has remained an open question how T cells outside the CNS detect and respond to microbial antigen expressed in the brain. The recent discovery of cerebrospinal fluid-draining lymphatic vessels in the dural meninges of mice and humans [12, 14] has challenged previous frameworks for how antigen exits the CNS [48–50]. Here, we provide evidence that meningeal lymphatic drainage supports dendritic cell activation and peripheral CD4^+^ and CD8^+^ T cell responses against the brain-tropic pathogen *T. gondii*. We further demonstrate that meningeal lymphatic drainage is dispensable for T cell responses against the parasite in the brain, despite reports showing drainage to be essential for the development of CNS-directed T cell responses in models of brain cancer [16, 44].

The deep cervical lymph nodes have been strongly implicated in mediating immune responses to CNS antigen [16, 17, 37]. Consistent with this notion, infection with the Pru-OVA strain of *T. gondii* led to robust expansion of SIINFEKL-specific T cells in the deep cervical lymph nodes, and the magnitude of this response tracked closely with parasite burden in the brain as infection progressed. In peripheral organs, foreign antigen is transported directly from the site of infection to lymph nodes for immune cell activation [51]. However, our data support a model in which antigen expressed in infected brain tissue must first reach the cerebrospinal fluid before it can be transported to lymph nodes for T cell activation. Indeed, parasite-specific T cell responses were significantly impaired in the DCLNs when lymphatic drainage of CSF components was restricted by surgical ligation. Future studies will be needed to more closely examine how CNS antigen is elaborated into the CSF. Most likely, this process is driven by “glymphatic” flow, a mechanism by which amyloid-β and other waste products have been shown to be cleared from the brain parenchyma [20, 52]. Glymphatic flow involves entry of CSF into the brain at Virchow-Robin spaces, exchange of macromolecules between the brain interstitial fluid and CSF across the glia limitans, and movement of this fluid back into the subarachnoid space via perivascular channels [19]. How *T. gondii* antigen makes it into the brain interstitium also needs to be addressed, and may depend on the release of antigen from infected host cells during the parasite’s lytic cycle of replication [53] or following pyroptotic host cell death [54]. Antigen might also be secreted by extracellular parasites after cyst reactivation or prior to invasion of new host cells.

In addition to the drainage of soluble antigen, lymphatic vessels promote the trafficking of dendritic cells from infected tissue after antigen uptake and phenotypic maturation [55]. Even though the dural meninges did not harbor active infection, we observed a significant accumulation of dendritic cells at this site, and these cells distributed specifically around the dural sinuses in regions where they could sample DQ-OVA injected into the CSF and access lymphatic vessels. These cells also expressed higher levels of co-stimulatory molecules in infected compared to naïve mice, although it remains to be determined what factors released in the brain, CSF, or meninges signal on these cells to induce maturation. Likely candidates include IFN-γ and TNF-α, which are highly expressed in the brain during infection [56], as well as pathogen-associated molecular patterns, including the TLR11 agonist *T. gondii* profilin [57], or damage-associated molecular patterns, which can accumulate in the CSF during *T. gondii* infection [58]. Notably, ligation of the lymphatic vessels reduced the number of dendritic cells in the deep cervical lymph nodes by nearly half and impaired dendritic cell maturation. These results suggest that infection of the brain promotes expansion and activation of dendritic cells at a distal anatomic site, one in which cells are optimally positioned to sample CNS-derived antigen and transport it to peripheral lymphoid tissues. By contrast, even though dendritic cells are observed to accumulate in the brain during *T. gondii* infection, it is unclear whether these cells are able to exit the CNS. Some studies have suggested that dendritic cells in the brain travel along the rostral migratory stream and exit by crawling along olfactory nerves [59, 60], but two-photon live imaging studies of *T. gondii*-infected mouse brains has revealed that CD11c+ cells, including tissue-resident macrophages and dendritic cells, are largely immotile [27]. EAE studies suggest that the function of dendritic cells in the CNS may be to support local reactivation of T cells or promote T cell entry into the CNS [61–63]. Thus, dendritic cells in the different anatomic compartments of the CNS likely play distinct roles in regulating immune responses to CNS antigen. Further studies are needed to clarify the function of dendritic cells at these sites, though challenges remain due to limitations in targeting cells within different compartments of the CNS.

Disrupting meningeal lymphatic drainage during chronic brain infection caused a striking impairment in parasite-specific T cell activation and proliferation in the DCLNs, but did not affect T cell responses in the brain. Because the T cell population in the brain requires continual replenishment by circulating T cells [7, 9], these results suggest that alternative sources of antigen likely exist outside the CNS-draining lymph nodes during chronic infection. In fact, we observed significant antigen-dependent proliferation of OT-I cells in both the deep cervical lymph nodes and the hindlimb-draining inguinal lymph nodes six weeks after infection. The source of antigen in the inguinal lymph nodes is still not completely clear. However, considering the robust T cell responses observed in the inguinal lymph nodes during acute infection, it is possible that residual antigen persisting after clearance of acute infection contributed to the proliferation of OT-I cells at this site during chronic infection. In models of influenza or vesicular stomatitis virus infection, antigen depots were found to persist for weeks to months after viral clearance [64, 65]. It is also possible that low-grade infection of skeletal muscle [66, 67] is responsible for the T cell responses observed in the inguinal lymph nodes. Studies examining T cell responses against bradyzoite-specific antigens could help clarify two important points: (1) the degree to which T cell activation in lymph nodes that drain non-CNS tissues is a result of chronic infection of these tissues and (2) the potential importance of CNS lymphatic drainage in generating immune responses to antigens specifically expressed by the latent form of the parasite.

The presence of peripheral antigen during *T. gondii* brain infection is not surprising, as most CNS pathogens display varying degrees of tropism for multiple tissues and originate at a peripheral site prior to hematogenous dissemination to the brain, CSF, or meninges [68]. In fact, peripheral infection may serve to “immunize” the host ahead of CNS infection, making the drainage of CNS antigen to the deep cervical lymph nodes more redundant. By contrast, when tumors originate within the CNS, meningeal lymphatic drainage was found to be indispensable for T cell-dependent immunity in the brain [16, 44], likely because there were not alternative sources of antigen outside the CNS. Consistent with this idea, when tumor cells were injected into the flank of mice at the same time as intracranial engraftment, survival benefits of VEGF-C-enhanced meningeal lymphatic drainage were abrogated [16]. These results harken back to early studies performed by Medawar in the 1940s showing that skin grafts in the brain were rejected more quickly and at higher rates when also engrafted to the chest of rabbits [69]. For these reasons, we predict that meningeal lymphatic drainage may not be necessary to generate T cell responses against metastatic brain tumors, given the presence of early peripheral antigen prior to seeding of the brain.

To conclude, our study provides novel insight into how T cells respond to antigen expressed in the brain by a neurotropic pathogen. Our data shed new light on how the dural meninges promotes neuroimmune communication, while refining our understanding of how meningeal lymphatic drainage helps to preserve the health of the CNS.

## METHODS

### Mice

All mice were housed at University of Virginia specific pathogen-free facilities with a 12-hour light/dark cycle. C57BL/6J (#000664), CBA/J (#000656), B6.Cg-Tg(Itgax-Venus)1Mnz/J (CD11c^YFP^ mice, #008829), C57BL/6-Tg(TcraTcrb)1100Mjb/J (OT-I mice, #003831), and C57BL/6-Tg(Nr4a1-EGFP/cre)820Khog/J (Nur77^GFP^ mice, #016617) strains were originally purchased from the Jackson Laboratory and then maintained within our animal facility. B6.SJL-*Ptprc^a^Pepc^b^*/BoyCrCrl (CD45.1 congenic mice, #564) and Swiss Webster (#024) strains were purchased from Charles River Laboratories. To generate OT-I (CD45.1 congenic) mice, OT-I and CD45.1 congenic strains were cross-bred within our facility. OT-I (CD45.1 congenic) mice were then crossed to Nur77^GFP^ mice to generate OT-I/Nur77^GFP^ (CD45.1 congenic) mice for use in adoptive transfer studies. For experiments reported in this study, age-matched young adult female mice (7-10 weeks of age) were used. All experiments were approved by the Institutional Animal Care and Use Committee at the University of Virginia under protocol number 3968.

### Parasite strains and infection

Mice were infected with avirulent, type II strains of *T. gondii*. The ME49 strain was maintained in chronically infected (2-6 months) Swiss Webster mice and passaged through CBA/J mice. For experimental infections with the ME49 strain, tissue cysts were prepared from homogenized brains of chronically infected (4-8 weeks) CBA/J mice. Mice were then inoculated intraperitoneally (i.p.) with 10 tissue cysts of ME49 in 200 μl of 1X PBS. The transgenic Prugniaud strain of *T. gondii* expressing ovalbumin (aa 140-386) and TdTomato (Pru-OVA) were generously provided by Anita Koshy (University of Arizona) and maintained by serial passage through human foreskin fibroblast (HFF) monolayers in parasite culture medium (DMEM [Gibco], 20% Medium 199 [Gibco], 10% FBS [Gibco], 1% penicillin/streptomycin [Gibco], and 10 μg/ml gentamicin [Gibco]). The parental Prugniaud strain (PruΔHPT, parental Pru) was similarly maintained. For experimental infections with the Pru-OVA or parental Pru strains, tachyzoites were purified from HFF cultures by needle-passaging scraped cells and filtering the parasites through a 5.0-μm filter (EMD Millipore). Mice were then inoculated i.p. with 1,000 tachyzoites of Pru-OVA or parental Pru in 200 μl of 1X PBS.

### Assessment of parasite burden

For assessment of parasite burden by real-time PCR, genomic DNA (gDNA) was isolated from mouse brains or dural meninges using the Isolate II Genomic DNA Kit (Bioline, BIO-52067) following the manufacturer’s instructions. Prior to tissue lysis, whole brains were mechanically homogenized using an Omni TH tissue homogenizer (Omni International). Amplification of the 529-bp repeat element in the *T. gondii* genome was performed as described previously [70], using the *Taq* polymerase-based SensiFAST™ Probe No-ROX Kit (Bioline, BIO-86005) and CFX384 Real-Time System (Bio-Rad) to assay 500 ng of DNA per sample. A standard curve generated from 10-fold serial dilutions of *T. gondii* gDNA, isolated from cultured HFFs and ranging from 3 to 300,000 genome copies, was used to determine the total number of *T. gondii* genome copies per tissue. For assessment of parasite burden by cyst counts, whole brains were minced with a razor blade and passed through an 18-gauge and 22-gauge needle to mechanically homogenize the tissue. 30 μl of tissue homogenate was then mounted on a microscope slide and *T. gondii* cysts were enumerated using a DM 2000 LED brightfield microscope (Leica).

### Intra-cisterna magna injections

Mice were anesthetized by i.p. injection of a solution containing ketamine (100 mg/kg) and xylazine (10 mg/kg) diluted in saline. Mice were secured in a stereotaxic frame with the head angled slightly downward. An incision in the skin was made at the base of the skull and the muscle layers overlying the atlanto-occipital membrane were retracted. A 33-gauge Hamilton syringe (#80308) was inserted at a steep angle through the membrane to inject the desired solution into the CSF-filled cisterna magna. After injection, the skin was closed using 5-0 nylon sutures and mice received a subcutaneous (s.c.) injection of ketoprofen (2 mg/kg). Mice were then allowed to recover on a heating pad until awake.

### Lymphatic vessel ligation

Collecting lymphatic vessels afferent to the deep cervical lymph nodes were surgically ligated as previously described [12]. Mice were anesthetized by i.p. injection of a solution containing ketamine (100 mg/kg) and xylazine (10 mg/kg) diluted in saline. A midline incision was made into the skin overlying the anterior neck after being shaved and cleaned with povidone-iodine and 70% ethanol. The sternocleidomastoid (SCM) muscles were retracted and collecting lymphatic vessels afferent to the DCLNs were ligated using 9-0 nylon suture (Living Systems Instrumentation, THR-G). After bilateral ligation of the lymphatic vessels, the skin was closed using 5-0 nylon sutures and mice received s.c. injection of ketoprofen (2 mg/kg). Mice were then allowed to recover on a heating pad until awake. Sham operations were performed in a similar manner, except lymphatic vessels were not ligated after retraction of the SCM muscles and exposure of the DCLNs.

### Cerebrospinal fluid tracer analysis

To assess the function of meningeal lymphatic drainage in mice that had received ligation surgery, tracers were injected into the cerebrospinal fluid (CSF) and outflow to the deep cervical lymph nodes was measured. For qualitative assessment, 5 μl of 10% Evans blue (Sigma-Aldrich) diluted in artificial CSF (aCSF, Harvard Apparatus) was introduced into the subarachnoid space of ligated or sham-operated mice by intra-cisterna magna (i.c.m.) injection. 1 h after injection, mice were sacrificed and drainage of the dye to the DCLNs was visualized under a S6D stereomicroscope (Leica). For quantitative assessment, 3 μl of Alexa Fluor 594-conjugated ovalbumin (OVA-AF594, Thermo Fisher) diluted in aCSF (2 mg/ml) was introduced into the subarachnoid space of ligated or sham-operated mice by i.c.m. injection. 2 h after injection, mice were sacrificed and the amount of OVA-AF594 present in the DCLNs was determined by fluorescence spectroscopy. Briefly, DCLNs were harvested and enzymatically digested with Collagenase D (1 mg/ml, Sigma-Aldrich) at 37°C for 30 min, followed by a 2 h treatment in T-PER™ Tissue Protein Extraction Reagent (Thermo Scientific) at 4°C. Samples were centrifuged at >10,000 rpm to remove cellular debris and the supernatant containing extracted protein was transferred to a black flat-bottom microwell plate (Millipore Sigma, M9685). Relative fluorescence intensity of each sample was measured on a SpectraMax iD3 microplate reader (Molecular Devices) using an excitation wavelength of 590 nm and emission wavelength of 630 nm and compared to a standard curve to determine the total amount of tracer that drained.

### DQ-OVA analysis

The uptake and processing of CSF-derived protein by antigen-presenting cells outside the CNS was evaluated by injecting 3 μl of DQ-OVA (2 mg/ml, Invitrogen) into the CSF of chronically infected C57BL/6 mice by i.c.m. injection. The fluorescent signal of proteolytically cleaved DQ-OVA, with a peak excitation wavelength of 505 nm and peak emission wavelength of 515 nm, was then detected by flow cytometry or confocal microscopy. For flow cytometry studies, the dural meninges, deep cervical lymph nodes, and inguinal lymph nodes were harvested and prepared as single-cell suspensions for cell surface staining 5 h after i.c.m. injection of DQ-OVA. For flow acquisition, fluorescence emitted by processed DQ-OVA was detected using the FL1 sensor (525/40 nm) of a Gallios flow cytometer (Beckman Coulter). For imaging studies, mice were sacrificed 12 h after injection of DQ-OVA, and whole-mount dural meninges were fixed, then immunostained using directly-conjugated antibodies targeting MHC class II (Super Bright™ 436, eBioscience) and LYVE1 (eFluor™ 660, eBioscience). Fluorescence emitted by processed DQ-OVA was detected using 488 nm laser excitation on a TCS SP8 confocal microscope (Leica).

### Immunohistochemistry

Image analysis was performed on dural tissue isolated from CD11c^YFP^ reporter mice and DQ-OVA-injected C57BL/6 mice. After sacrifice, transcardiac perfusion of mice was performed using 20 ml of cold 1X PBS. The dorsal aspect of the skull was removed and the dural meninges were fixed, while still attached to the skull bone, in 4% PFA for 6-8 h at 4°C. Then, as previously described [71], whole-mount dural meninges were carefully dissected, washed, and incubated in a blocking solution (2% normal donkey serum, 1% BSA, 0.05% Tween 20, and 0.5% Triton X-100 in 1X PBS) at room temperature for 45 min. Next, tissue was stained for 2 h at room temperature with directly-conjugated primary antibodies (1:100 dilution). Tissue was then washed three times in a solution of 0.05% Tween 20 (in 1X PBS), mounted onto glass slides using AquaMount (Fisher Scientific), and coverslipped. In some experiments, the tissue was counter-stained with DAPI (Thermo Scientific) and washed just before mounting onto slides. Primary antibodies (from eBioscience) included: MHC class II (I-A/I-E)-Super Bright 436 (62- 5321-80), LYVE1-EF570 (41-0443-82), and LYVE1-EF660 (50-0443-82). Images were acquired using a Leica TCS SP8 confocal microscope and analyzed using Fiji software [72]. Experimenter was blinded to the identity of experimental group during image analysis.

### Tissue processing for flow cytometry

After sacrifice, transcardiac perfusion of mice was performed using 20 ml of cold 1X PBS. For analysis of the dural meninges, the dorsal aspect of the skull was removed and the dural meninges were carefully dissected from the skull bone under a S6D stereomicroscope (Leica). The tissue was then digested in 1X HBSS (without Ca^2+^ or Mg^2+^, Gibco) with Collagenase D (1 mg/ml, Sigma-Aldrich) and Collagenase VIII (1 mg/ml, Sigma-Aldrich) at 37°C for 30 min, before being passed through a 70-μm strainer (Corning). Cells were then pelleted, resuspended, and kept on ice. Deep cervical lymph nodes (DCLNs), which are positioned laterally to the trachea and underneath the sternocleidomastoid muscles, and inguinal lymph nodes (ILNs) were harvested bilaterally into cold complete RPMI media (cRPMI; 10% FBS [Gibco], 1% penicillin/streptomycin [Gibco], 1% sodium pyruvate [Gibco], 1% non-essential amino acids [Gibco], and 0.1% 2-Mercaptoethanol [Life Technologies]). Lymph nodes were mechanically homogenized and gently pressed through a 70-μm strainer (Corning). Cells were then pelleted, resuspended, and kept on ice. Brains were harvested into cold cRPMI, minced with a razor blade, and passed through an 18-gauge and 22-gauge needle for mechanical homogenization. Tissue was then digested in a solution containing collagenase/dispase (0.227 mg/ml, Sigma-Aldrich) and DNase (50 U/ml, Roche) at 37 °C for 45-60 min, before being passed through a 70-μm strainer (Corning) and washed with cRPMI. Myelin was separated from mononuclear cells by resuspending samples in 20 ml of 40% Percoll (Cytiva) and centrifuging at 650*g* for 25 min. Myelin was aspirated and cell pellets were washed, resuspended, and kept on ice. Spleens were harvested into cold cRPMI, then mechanically homogenized and washed through a 40-μm strainer (Corning). Cells were resuspended in RBC lysis solution (0.16 M NH_4_Cl) for 2 min. Cells were then washed, resuspended, and kept on ice.

### Cerebrospinal fluid collection for flow cytometry

Mice were anesthetized by i.p. injection of a solution containing ketamine (100 mg/kg) and xylazine (10 mg/kg) diluted in saline. An incision in the skin was made at the base of the skull and the muscle layers overlying the atlanto-occipital membrane were retracted. A pulled glass capillary (Sutter Instrument, BF100-50-10) was inserted through the membrane into the cisterna magna and 5-10 μl of CSF was collected per mouse. CSF from 4-5 mice was pooled, and a total of 25 μl of CSF per pooled sample was analyzed by flow cytometry.

### Flow cytometry

Single-cell suspensions prepared from tissues or cerebrospinal fluid were pipetted into a 96-well plate and pelleted. Cells were treated with 50 μl of Fc block (0.1% rat gamma globulin [Jackson ImmunoResearch], 1 μg/ml of 2.4G2 [BioXCell]) for 10 min at room temperature. Cells were then stained for surface markers and incubated with a fixable live/dead viability dye for 30 min at 4°C. In experiments where SIINFEKL-specific CD8+ T cells were analyzed, cells were pre-incubated with PE-conjugated H-2K^b^/OVA (SIINFEKL) tetramer (NIH Tetramer Core Facility) for 15 min at room temperature prior to cell surface staining. Splenocytes were only stained with the live/dead viability dye and were used as a compensation control. After surface staining, cells were washed with FACS buffer (0.2% BSA and 2 mM EDTA in 1X PBS) and, if intracellular staining was performed, cells were treated with a fixation/permeabilization solution (eBioscience, 00-5123-43 and 00-5223-56) overnight then stained for intracellular markers in permeabilization buffer (eBioscience, 00-8333-56) for 30 min at 4°C. Finally, samples were resuspended in FACS buffer and acquired on a Gallios flow cytometer (Beckman Coulter). Samples were analyzed using FlowJo software v.10. For dural meninges and cerebrospinal fluid, cell counts were determined using absolute counting beads (Life Technologies, C36950) pipetted into samples just prior to acquisition. For lymph nodes and brain, cell counts were measured using a Hausser Scientific™ hemacytometer (Fisher Scientific). Cells were stained for surface markers using the following eBioscience antibodies at 1:200 dilution: CD45-FITC (11-0451-82), CD62L- FITC (11-0621-85), CD80-FITC (11-0801-82), MHC class II (I-A/I-E)-PE (12-5321-82), CD69- PE (12-0691-82), CD11c-PerCP-Cy5.5 (45-0114-82), CD4-PerCP-Cy5.5 (45-0042-82), CD45-PerCP-Cy5.5 (45-0451-82), CD11b-PerCP-Cy5.5 (45-0112-82), CD8α-PerCP-Cy5.5 (45-0081- 82), CD45.1-PerCP-Cy5.5 (45-0453-82), CD80-PE-Cy7 (25-0801-82), CD11c-PE-Cy7 (25-0114-82), CD4-PE-Cy7 (25-0041-82), CD8α-PE-Cy7 (25-0081-82), TCR-β-APC (17-5961-82), NK1.1-APC (17-5941-82), CD19-APC (17-0193-82), CD44-AF780 (47-0441-82), CD11b-AF780 (47-0112-82), CD86-EF450 (48-0862-82), MHC class II (I-A/I-E)-EF450 (48-5321-82), CD69-EF450 (48-0691-82), and CD4-EF450 (48-0042-82). Cells were stained for intracellular markers using the following eBioscience antibodies at 1:200 dilution: TNF-α-AF488 (53-7321- 82), IFN-γ-PerCP-Cy5.5 (45-7311-82), and Ki67-PE-Cy7 (25-5698-82). The following eBioscience live/dead dyes were used at a 1:800 dilution: Fixable Viability Dye eFluor 780 (65- 0865-14) and Fixable Viability Dye eFluor 506 (65-0866-18).

### *Ex vivo* T cell restimulation

Cells isolated from the deep cervical lymph nodes or brain were seeded at 2.5 x 10^5^ cells per well and restimulated for 24 h with soluble tachyzoite antigen (STAg) at 25 μg/ml. Cells were then incubated with brefeldin A (10 μg/ml, Selleck Chemicals) at 37°C for 6 h. Cells were washed and stained for surface markers, and then intracellular cytokine staining was performed. To prepare STAg, tachyzoites of the RH strain of *T. gondii* were purified from HFF cultures, filtered through a 5.0-μm filter (EMD Millipore), resuspended in 1X PBS, and lysed by five freeze-thaw cycles. Protein concentration was determined by BCA assay (Pierce) and stock solutions were stored at −80°C.

### OT-I cell labeling and transfer

Spleens and lymph nodes from OT-I/Nur77-GFP (CD45.1 congenic) mice were pooled, mechanically dissociated, treated with RBC lysis solution (0.16 M NH_4_Cl), and passed through a 70-μm strainer (Corning). Then, OT-I cells were expanded *in vitro* as previously described [7]. Briefly, isolated cells were cultured for 24 h with ovalbumin protein (Worthington, LS003048) at 500 μg/ml. After 24 h, cells were washed and rested. On days 4 and 6, cells were treated with recombinant IL-2 (Proleukin, Prometheus Laboratories) at 200 U/ml. On day 7, cells were washed and labeled with 10 μM CellTrace Violet (Invitrogen, C34557) for 20 min at 37°C. Cells were resuspended in 1X PBS and then 4×10^6^ to 5×10^6^ labeled OT-I cells were transferred to mice anesthetized with ketamine (100 mg/kg) and xylazine (10 mg/kg) by retro-orbital intravenous injection.

### Statistical analysis

Statistical analyses were performed using Prism software (v8.4) or RStudio (v1.1) statistical packages. Power analysis was performed in R using the pwr software package to calculate group sizes needed to achieve a power of 0.8, with effect size estimated from preliminary experiments. Randomization of samples was ensured by including mice from the same experimental group in different cages. A two-tailed student’s *t*-test was used to compare two independent groups. A one-way ANOVA was performed to compare three or more independent groups, with correction for multiple testing by Tukey’s procedure. When data were pooled from multiple experiments, a randomized block ANOVA was performed using the lme4 software package in R [73]. This test models experimental groups as a fixed effect and experimental day as a random effect. The test used for each experiment is denoted in the figure legend, and *p* values are similarly indicated, with ns = not significant, *p* < 0.05 (*), *p* < 0.01 (**), and *p* < 0.001 (***). Graphs were generated using Prism software and show mean values ± s.e.m. along with individual data points representative of individual mice (biological replicates).

## ACKNOWLEDGMENTS

We would like to acknowledge members of the Center for Brain Immunology and Glia (BIG) at the University of Virginia for their scientific input, training in surgical techniques, and access to instrumentation as this project was being developed. Schematic diagrams were generated using BioRender (https://biorender.com/). We thank Marieke K. Jones for her guidance with statistical analyses and R programming. We thank Anita Koshy at the University of Arizona for providing transgenic parasite strains used in this study. We also acknowledge the support we received from the Biomolecular Analysis Facility at the University of Virginia.

## AUTHOR CONTRIBUTIONS

M.A.K. designed and performed experiments, analyzed data, and wrote the manuscript. M.N.C., I.W.B., L.A.S., K.S., and S.J.B. assisted with experimental procedures and provided input on experimental design. S.A.L. and I.S. assisted with experimental procedures. T.H.H. supervised the work and edited the manuscript.

---

**Figure 1 – Supplementary 1.**
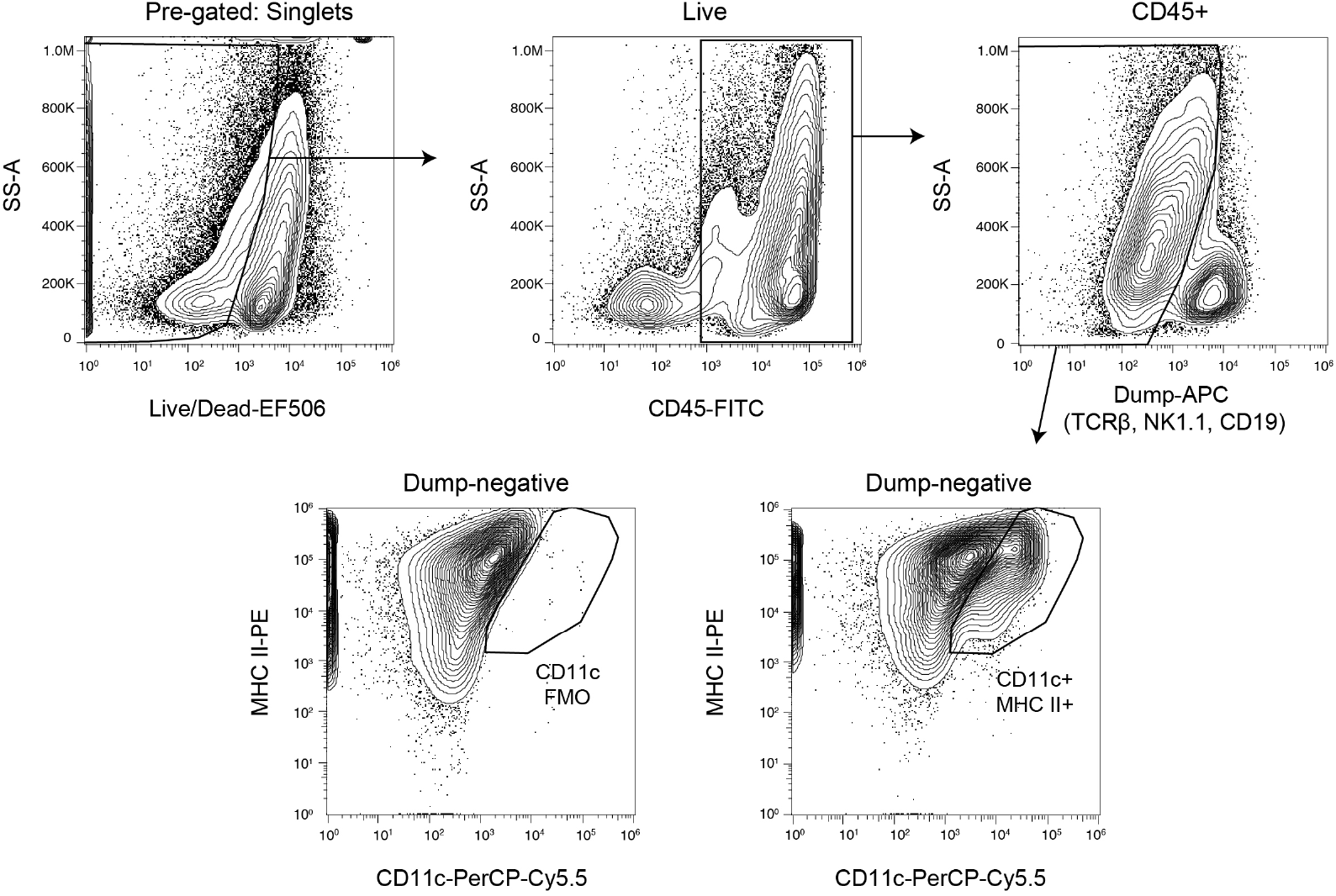
Gating strategy for dural dendritic cells. Gating strategy for identification of live dendritic cells (CD45^+^TCRβ^-^NK1.1^-^CD19^-^CD11c^hi^MHC II^hi^) in single-cell suspensions of enzymatically digested dural meninges. Pre-gated on singlets.

**Figure 1 – Supplementary 2.**
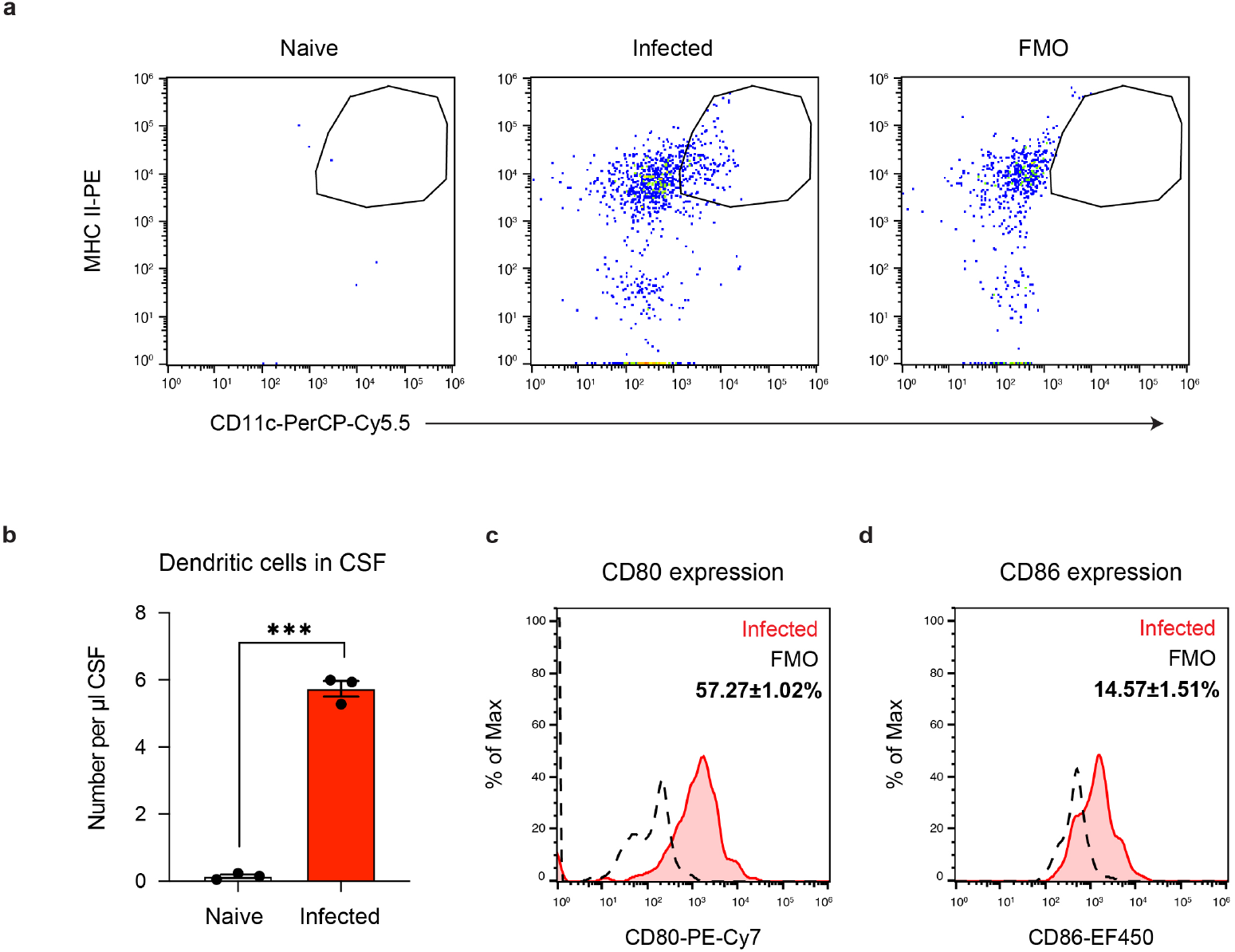
Dendritic cells are present in the cerebrospinal fluid of infected but not uninfected mice. Cerebrospinal fluid (CSF) was isolated and pooled from 4-5 naïve or chronically infected C57BL/6 mice at 6 wpi. Surface staining was performed to identify dendritic cells by flow cytometry. **a**, Representative dot plots of CD11c^hi^MHC II^hi^ dendritic cells pre-gated on singlets/live/CD45^+^/TCRβ^-^/NK1.1^-^/CD19^-^. **b**, Quantification of dendritic cell number per μl of CSF in naïve and chronically infected mice. Data are compiled from three experiments and are represented as mean values ± s.e.m. Each data point represents the concentration of cells found in the pooled CSF of 4-5 mice. Statistical significance was measured using a two-tailed unpaired student’s *t*-test, with *p* < 0.001 (***). **c-d**, Representative flow histograms showing CD80 **(c)** and CD86 **(d)** expression by dendritic cells in the CSF of chronically infected mice. The average frequency of CD80^+^ or CD86^+^ dendritic cells (mean value ± s.e.m.) was calculated from three pooled experiments.

**Figure 2 – Supplementary 1.**
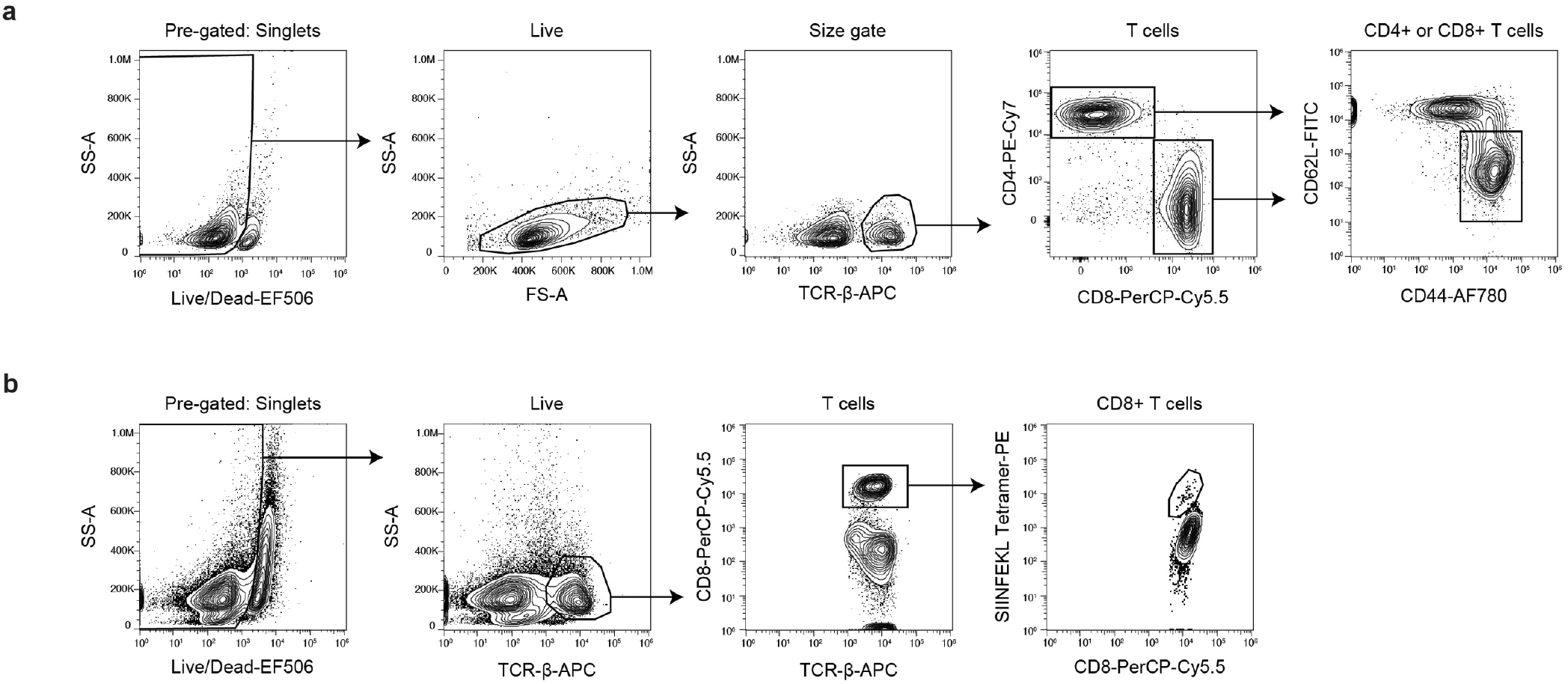
Gating strategy for CD4^+^ and CD8^+^ T cell responses following infection with ME49 or Pru-OVA. **a**, Gating strategy for identification of CD44^hi^CD62L^lo^ (activated) CD4^+^ or CD8^+^ T cells in lymph nodes of mice infected with ME49. Pre-gated on singlets. **b,** Gating strategy for identification of SIINFEKL-specific CD8^+^ T cells in lymph nodes of mice infected with Pru-OVA. Pre-gated on singlets.

**Figure 3 – Supplementary 1:**
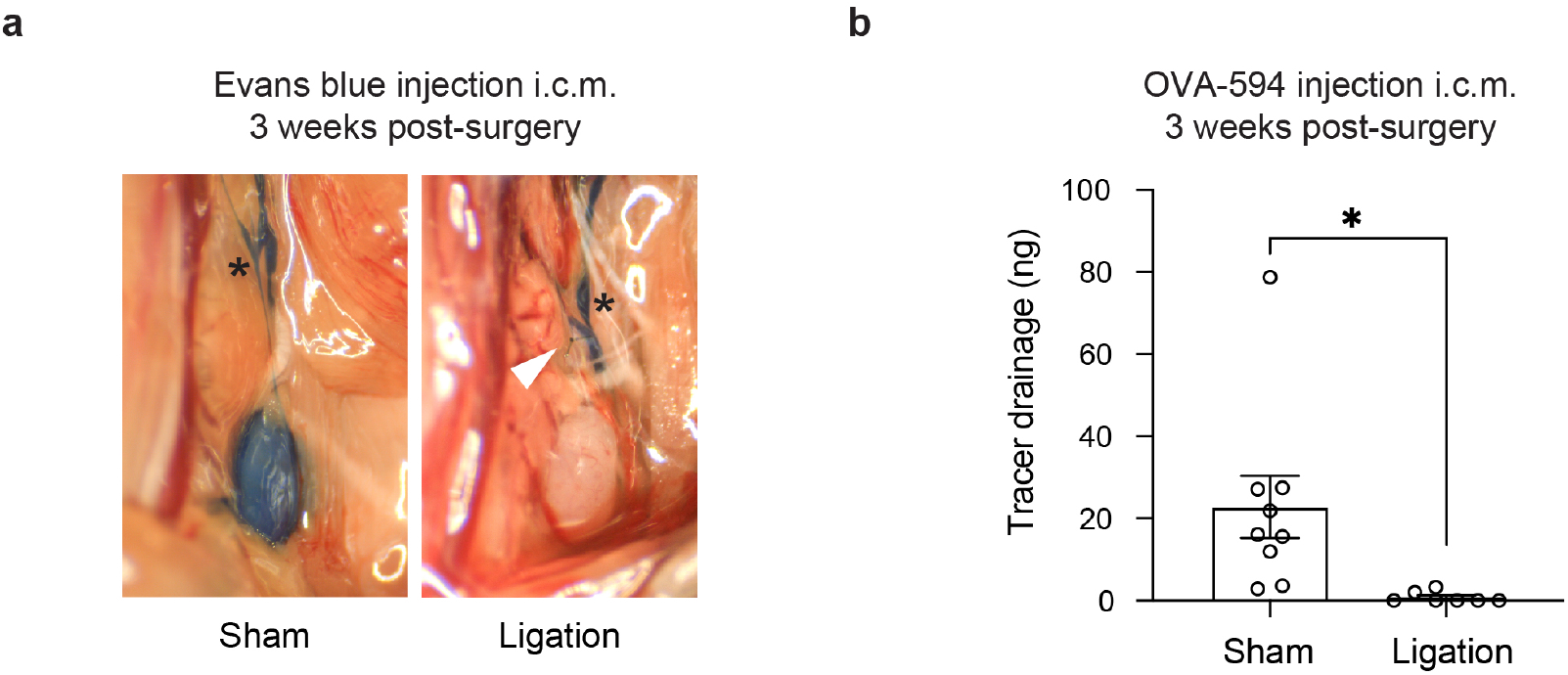
Tracer studies confirm disruption of meningeal lymphatic drainage by ligation surgery. Chronically infected C57BL/6 mice were subjected to surgical ligation of collecting vessels afferent to the DCLNs or sham surgery. **a**, Representative image of Evans blue dye drainage to the DCLNs 1 h after i.c.m. injection of ligated or sham-operated mice. Suture placement is indicated by a white arrow (right). CSF-derived dye flows through collecting lymphatic vessels of ligated and sham-operated mice (asterisks) but only reaches the DCLNs of sham-operated mice (left). Two independent experiments were performed. **b**, Quantification of OVA-AF594 protein tracer accumulation in DCLN lysates of ligated or sham-operated mice 2 h after i.c.m. injection by fluorescence spectroscopy. Data are compiled from two experiments and are represented as mean values ± s.e.m. Statistical significance was measured using randomized block ANOVA (two-way), with *p* < 0.05 (*).

**Figure 3 – Supplementary 2:**
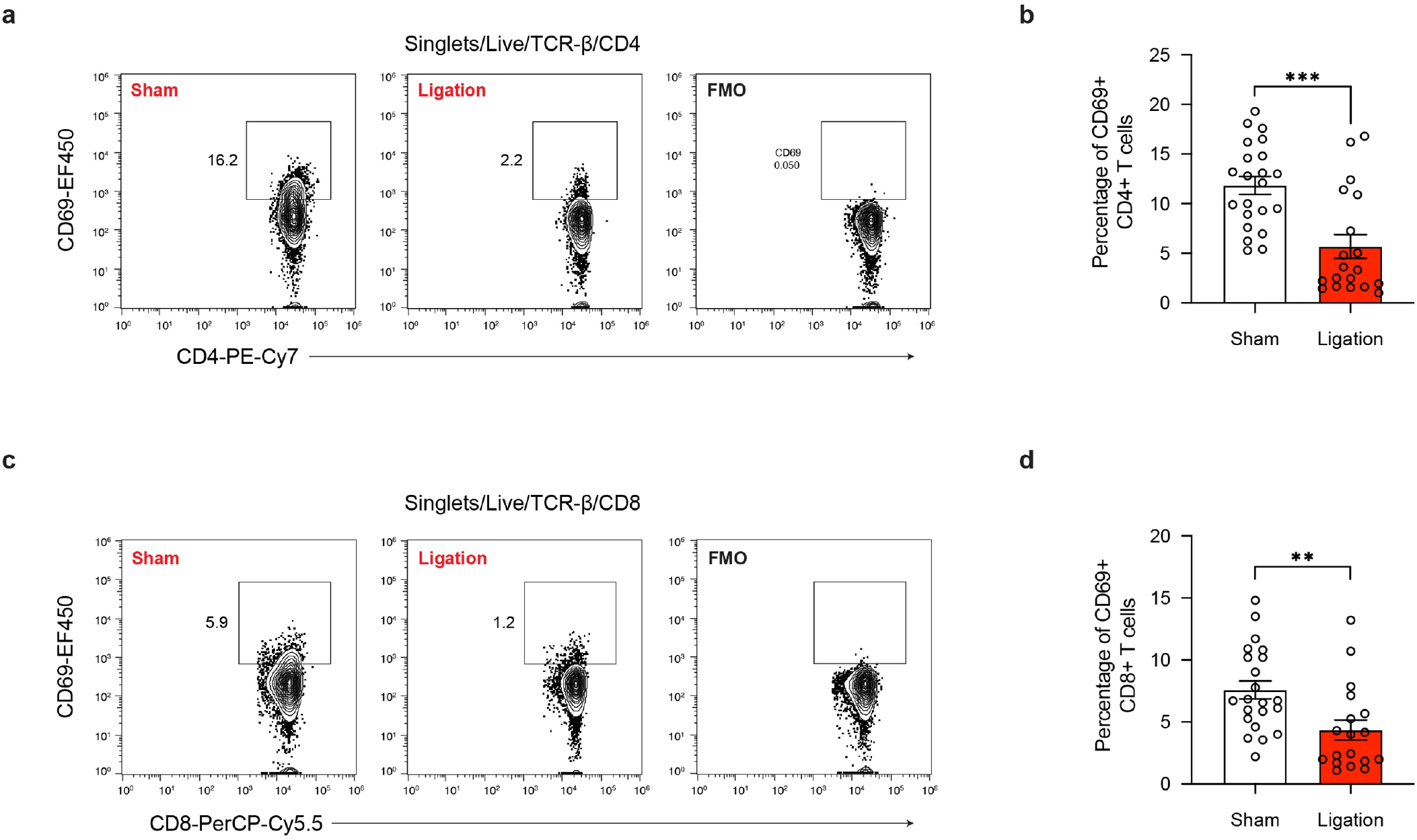
Restricting meningeal lymphatic drainage impairs T cell activation in the deep cervical lymph nodes. **a-b**, Representative contour plots **(a)** and average frequency **(b)** of CD69-expressing CD4^+^ T cells in the DCLNs of ligated or sham-operated mice. **c-d**, Representative contour plots **(c)** and average frequency **(d)** of CD69-expressing CD8^+^ T cells in the DCLNs of ligated or sham-operated mice. Data are compiled from four experiments and are represented as mean values ± s.e.m. Statistical significance was measured using randomized block ANOVA (two-way), with *p* < 0.01 (**) and *p* < 0.001 (***).

**Figure 3 – Supplementary 3:**
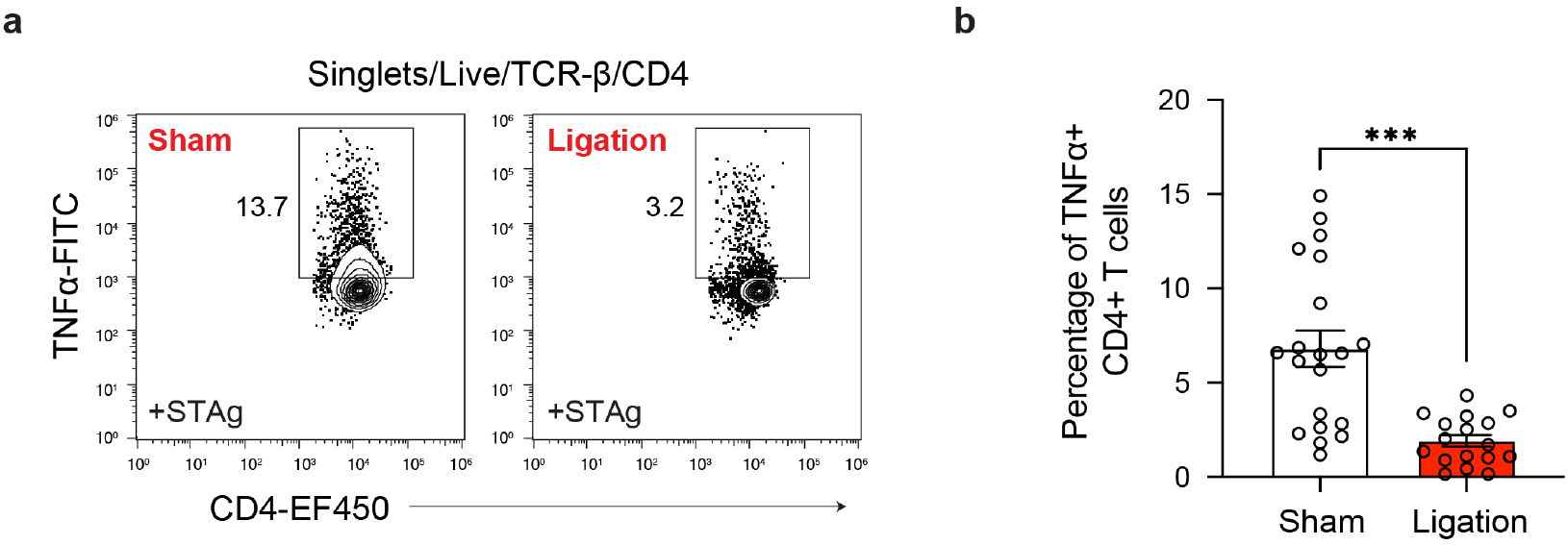
Restricting meningeal lymphatic drainage impairs TNF-α production in the deep cervical lymph nodes. **a-b**, Intracellular staining of TNF-α was performed on cells isolated from the DCLNs of ligated or sham-operated mice 24 h after *ex vivo* restimulation with soluble tachyzoite antigen (STAg). Representative contour plots **(a)** and average frequency **(b)** of CD4^+^ T cells expressing TNF-α in response to STAg are given. Data are compiled from four experiments and are represented as mean values ± s.e.m. Statistical significance was measured using randomized block ANOVA (two-way), with *p* < 0.001 (***).

**Figure 5 – Supplement 1:**
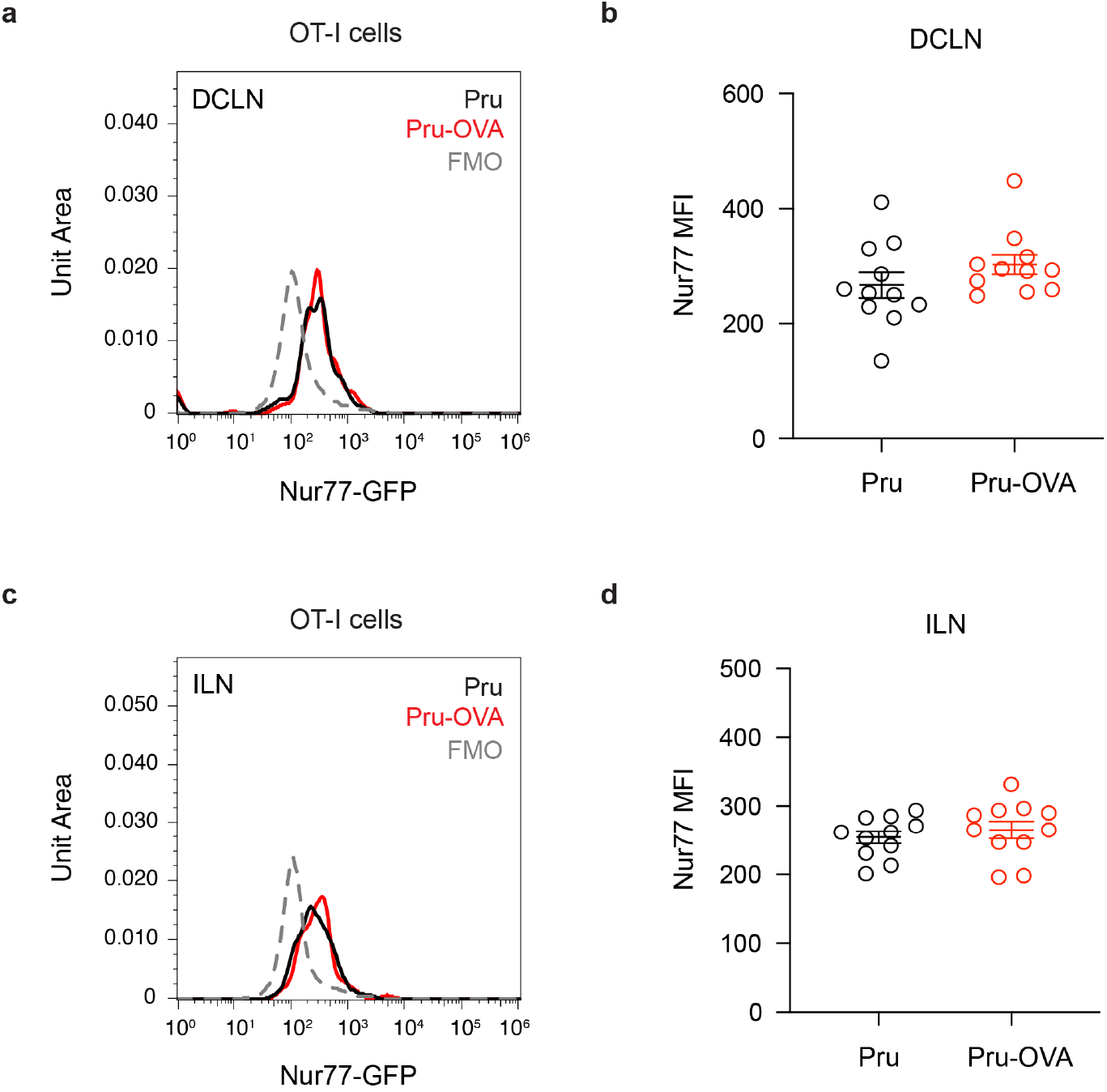
Assessing Nur77-GFP expression in mice chronically infected with Pru-OVA or parental Pru. Nur77^GFP^ expression was measured seven days after adoptive transfer of OT-I/Nur77^GFP^ cells (CD45.1 congenic) to C57BL/6 mice chronically infected with Pru-OVA or parental Pru. **a-b**, Representative flow histogram **(a)** and average geometric mean fluorescence intensity (MFI) **(b)** of Nur77-GFP expressed by OT-I cells of the deep cervical lymph nodes (DCLN). **c-d**, Representative flow histogram **(c)** and average MFI **(d)** of Nur77-GFP expressed by OT-I cells of the inguinal lymph nodes (ILN).

## REFERENCES

1. Ellwardt, E., et al., Understanding the Role of T Cells in CNS Homeostasis. Trends Immunol, 2016. 37(2): p. 154–165.

2. Korn, T. and A. Kallies, T cell responses in the central nervous system. Nat Rev Immunol, 2017. 17(3): p. 179–194.

3. Hill, D. and J.P. Dubey, Toxoplasma gondii: transmission, diagnosis and prevention. Clin Microbiol Infect, 2002. 8(10): p. 634–40.

4. Pappas, G., N. Roussos, and M.E. Falagas, Toxoplasmosis snapshots: global status of Toxoplasma gondii seroprevalence and implications for pregnancy and congenital toxoplasmosis. Int J Parasitol, 2009. 39(12): p. 1385–94.

5. Luft, B.J. and J.S. Remington, Toxoplasmic encephalitis in AIDS. Clin Infect Dis, 1992. 15(2): p. 211–22.

6. Gazzinelli, R., et al., Simultaneous depletion of CD4+ and CD8+ T lymphocytes is required to reactivate chronic infection with Toxoplasma gondii. J Immunol, 1992. 149(1): p. 175–80.

7. Harris, T.H., et al., Generalized Levy walks and the role of chemokines in migration of effector CD8+ T cells. Nature, 2012. 486(7404): p. 545–8.

8. Schaeffer, M., et al., Dynamic imaging of T cell-parasite interactions in the brains of mice chronically infected with Toxoplasma gondii. J Immunol, 2009. 182(10): p. 6379–93.

9. Wilson, E.H., et al., Behavior of parasite-specific effector CD8+ T cells in the brain and visualization of a kinesis-associated system of reticular fibers. Immunity, 2009. 30(2): p. 300–11.

10. Thomas, S.N., N.A. Rohner, and E.E. Edwards, Implications of Lymphatic Transport to Lymph Nodes in Immunity and Immunotherapy. Annu Rev Biomed Eng, 2016. 18: p. 207–33.

11. Gasteiger, G., M. Ataide, and W. Kastenmuller, Lymph node - an organ for T-cell activation and pathogen defense. Immunol Rev, 2016. 271(1): p. 200–20.

12. Louveau, A., et al., Structural and functional features of central nervous system lymphatic vessels. Nature, 2015. 523(7560): p. 337–41.

13. Aspelund, A., et al., A dural lymphatic vascular system that drains brain interstitial fluid and macromolecules. J Exp Med, 2015. 212(7): p. 991–9.

14. Absinta, M., et al., Human and nonhuman primate meninges harbor lymphatic vessels that can be visualized noninvasively by MRI. Elife, 2017. 6.

15. Albayram, M.S., et al., Non-invasive MR imaging of human brain lymphatic networks with connections to cervical lymph nodes. Nat Commun, 2022. 13(1): p. 203.

16. Song, E., et al., VEGF-C-driven lymphatic drainage enables immunosurveillance of brain tumours. Nature, 2020. 577(7792): p. 689–694.

17. Louveau, A., et al., CNS lymphatic drainage and neuroinflammation are regulated by meningeal lymphatic vasculature. Nat Neurosci, 2018. 21(10): p. 1380–1391.

18. Abbott, N.J., Evidence for bulk flow of brain interstitial fluid: significance for physiology and pathology. Neurochem Int, 2004. 45(4): p. 545–52.

19. Plog, B.A. and M. Nedergaard, The Glymphatic System in Central Nervous System Health and Disease: Past, Present, and Future. Annu Rev Pathol, 2018. 13: p. 379–394.

20. Iliff, J.J., et al., A paravascular pathway facilitates CSF flow through the brain parenchyma and the clearance of interstitial solutes, including amyloid beta. Sci Transl Med, 2012. 4(147): p. 147ra111.

21. Ling, C., M. Sandor, and Z. Fabry, In situ processing and distribution of intracerebrally injected OVA in the CNS. J Neuroimmunol, 2003. 141(1-2): p. 90–8.

22. Harris, M.G., et al., Immune privilege of the CNS is not the consequence of limited antigen sampling. Sci Rep, 2014. 4: p. 4422.

23. Li, X., et al., Meningeal lymphatic vessels mediate neurotropic viral drainage from the central nervous system. Nat Neurosci, 2022. 25(5): p. 577–587.

24. Alves de Lima, K., J. Rustenhoven, and J. Kipnis, Meningeal Immunity and Its Function in Maintenance of the Central Nervous System in Health and Disease. Annu Rev Immunol, 2020. 38: p. 597–620.

25. Rua, R. and D.B. McGavern, Advances in Meningeal Immunity. Trends Mol Med, 2018. 24(6): p. 542–559.

26. Rustenhoven, J., et al., Functional characterization of the dural sinuses as a neuroimmune interface. Cell, 2021. 184(4): p. 1000–1016 e27.

27. John, B., et al., Analysis of behavior and trafficking of dendritic cells within the brain during toxoplasmic encephalitis. PLoS Pathog, 2011. 7(9): p. e1002246.

28. Fischer, H.G., U. Bonifas, and G. Reichmann, Phenotype and functions of brain dendritic cells emerging during chronic infection of mice with Toxoplasma gondii. J Immunol, 2000. 164(9): p. 4826–34.

29. Carare, R.O., et al., Solutes, but not cells, drain from the brain parenchyma along basement membranes of capillaries and arteries: significance for cerebral amyloid angiopathy and neuroimmunology. Neuropathol Appl Neurobiol, 2008. 34(2): p. 131–44.

30. Clarkson, B.D., et al., CCR7 deficient inflammatory Dendritic Cells are retained in the Central Nervous System. Sci Rep, 2017. 7: p. 42856.

31. Cugurra, A., et al., Skull and vertebral bone marrow are myeloid cell reservoirs for the meninges and CNS parenchyma. Science, 2021. 373(6553).

32. Pulous, F.E., et al., Cerebrospinal fluid can exit into the skull bone marrow and instruct cranial hematopoiesis in mice with bacterial meningitis. Nat Neurosci, 2022. 25(5): p. 567–576.

33. Hammer, G.E. and A. Ma, Molecular control of steady-state dendritic cell maturation and immune homeostasis. Annu Rev Immunol, 2013. 31: p. 743–91.

34. Pashenkov, M., et al., Two subsets of dendritic cells are present in human cerebrospinal fluid. Brain, 2001. 124(Pt 3): p. 480–92.

35. Gerner, M.Y., et al., Dendritic cell and antigen dispersal landscapes regulate T cell immunity. J Exp Med, 2017. 214(10): p. 3105–3122.

36. Sixt, M., et al., The conduit system transports soluble antigens from the afferent lymph to resident dendritic cells in the T cell area of the lymph node. Immunity, 2005. 22(1): p. 19–29.

37. Harling-Berg, C., et al., Role of cervical lymph nodes in the systemic humoral immune response to human serum albumin microinfused into rat cerebrospinal fluid. J Neuroimmunol, 1989. 25(2-3): p. 185–93.

38. van Zwam, M., et al., Surgical excision of CNS-draining lymph nodes reduces relapse severity in chronic-relapsing experimental autoimmune encephalomyelitis. J Pathol, 2009. 217(4): p. 543–51.

39. Furtado, G.C., et al., Swift entry of myelin-specific T lymphocytes into the central nervous system in spontaneous autoimmune encephalomyelitis. J Immunol, 2008. 181(7): p. 4648–55.

40. Saeij, J.P., et al., Bioluminescence imaging of Toxoplasma gondii infection in living mice reveals dramatic differences between strains. Infect Immun, 2005. 73(2): p. 695–702.

41. Pepper, M., et al., Development of a system to study CD4+-T-cell responses to transgenic ovalbumin-expressing Toxoplasma gondii during toxoplasmosis. Infect Immun, 2004. 72(12): p. 7240–6.

42. Schluter, D., et al., Both lymphotoxin-alpha and TNF are crucial for control of Toxoplasma gondii in the central nervous system. J Immunol, 2003. 170(12): p. 6172–82.

43. Wang, X., et al., Gamma interferon production, but not perforin-mediated cytolytic activity, of T cells is required for prevention of toxoplasmic encephalitis in BALB/c mice genetically resistant to the disease. Infect Immun, 2004. 72(8): p. 4432–8.

44. Hu, X., et al., Meningeal lymphatic vessels regulate brain tumor drainage and immunity. Cell Res, 2020. 30(3): p. 229–243.

45. Moran, A.E., et al., T cell receptor signal strength in Treg and iNKT cell development demonstrated by a novel fluorescent reporter mouse. J Exp Med, 2011. 208(6): p. 1279–89.

46. Ashouri, J.F. and A. Weiss, Endogenous Nur77 Is a Specific Indicator of Antigen Receptor Signaling in Human T and B Cells. J Immunol, 2017. 198(2): p. 657–668.

47. Louveau, A., T.H. Harris, and J. Kipnis, Revisiting the Mechanisms of CNS Immune Privilege. Trends Immunol, 2015. 36(10): p. 569–577.

48. Walter, B.A., et al., The olfactory route for cerebrospinal fluid drainage into the peripheral lymphatic system. Neuropathol Appl Neurobiol, 2006. 32(4): p. 388–96.

49. Goldmann, J., et al., T cells traffic from brain to cervical lymph nodes via the cribroid plate and the nasal mucosa. J Leukoc Biol, 2006. 80(4): p. 797–801.

50. Weller, R.O., et al., Pathophysiology of the lymphatic drainage of the central nervous system: Implications for pathogenesis and therapy of multiple sclerosis. Pathophysiology, 2010. 17(4): p. 295–306.

51. Liao, S. and P.Y. von der Weid, Lymphatic system: an active pathway for immune protection. Semin Cell Dev Biol, 2015. 38: p. 83–9.

52. Iliff, J.J., et al., Impairment of glymphatic pathway function promotes tau pathology after traumatic brain injury. J Neurosci, 2014. 34(49): p. 16180–93.

53. Blader, I.J., et al., Lytic Cycle of Toxoplasma gondii: 15 Years Later. Annu Rev Microbiol, 2015. 69: p. 463–85.

54. Batista, S.J., et al., Gasdermin-D-dependent IL-1alpha release from microglia promotes protective immunity during chronic Toxoplasma gondii infection. Nat Commun, 2020. 11(1): p. 3687.

55. Russo, E., M. Nitschke, and C. Halin, Dendritic cell interactions with lymphatic endothelium. Lymphat Res Biol, 2013. 11(3): p. 172–82.

56. Matta, S.K., et al., Toxoplasma gondii infection and its implications within the central nervous system. Nat Rev Microbiol, 2021. 19(7): p. 467–480.

57. Yarovinsky, F., et al., TLR11 activation of dendritic cells by a protozoan profilin-like protein. Science, 2005. 308(5728): p. 1626–9.

58. Still, K.M., et al., Astrocytes promote a protective immune response to brain Toxoplasma gondii infection via IL-33-ST2 signaling. PLoS Pathog, 2020. 16(10): p. e1009027.

59. Mohammad, M.G., et al., Immune cell trafficking from the brain maintains CNS immune tolerance. J Clin Invest, 2014. 124(3): p. 1228–41.

60. Kaminski, M., et al., Migration of monocytes after intracerebral injection at entorhinal cortex lesion site. J Leukoc Biol, 2012. 92(1): p. 31–9.

61. Goverman, J., Autoimmune T cell responses in the central nervous system. Nat Rev Immunol, 2009. 9(6): p. 393–407.

62. D’Agostino, P.M., et al., Brain dendritic cells: biology and pathology. Acta Neuropathol, 2012. 124(5): p. 599–614.

63. Mundt, S., et al., Conventional DCs sample and present myelin antigens in the healthy CNS and allow parenchymal T cell entry to initiate neuroinflammation. Sci Immunol, 2019. 4(31).

64. Turner, D.L., et al., Persistent antigen presentation after acute vesicular stomatitis virus infection. J Virol, 2007. 81(4): p. 2039–46.

65. Zammit, D.J., et al., Residual antigen presentation after influenza virus infection affects CD8 T cell activation and migration. Immunity, 2006. 24(4): p. 439–49.

66. Jin, R.M., et al., Regulatory T Cells Promote Myositis and Muscle Damage in Toxoplasma gondii Infection. J Immunol, 2017. 198(1): p. 352–362.

67. Melchor, S.J., et al., T. gondii infection induces IL-1R dependent chronic cachexia and perivascular fibrosis in the liver and skeletal muscle. Sci Rep, 2020. 10(1): p. 15724.

68. Cain, M.D., et al., Mechanisms of Pathogen Invasion into the Central Nervous System. Neuron, 2019. 103(5): p. 771–783.

69. Medawar, P.B., Immunity to homologous grafted skin; the fate of skin homografts transplanted to the brain, to subcutaneous tissue, and to the anterior chamber of the eye. Br J Exp Pathol, 1948. 29(1): p. 58–69.

70. Homan, W.L., et al., Identification of a 200- to 300-fold repetitive 529 bp DNA fragment in Toxoplasma gondii, and its use for diagnostic and quantitative PCR. Int J Parasitol, 2000. 30(1): p. 69–75.

71. Louveau, A., A.J. Filiano, and J. Kipnis, Meningeal whole mount preparation and characterization of neural cells by flow cytometry. Curr Protoc Immunol, 2018. 121(1): p. e50.

72. Schindelin, J., et al., Fiji: an open-source platform for biological-image analysis. Nat Methods, 2012. 9(7): p. 676–82.

73. Bates, D., et al., Fitting Linear Mixed-Effects Models Using lme4. Journal of Statistical Software, 2015. 67(1): p. 1–48.

